# Antagonistic pleiotropy plays an important role in governing the evolution and genetic diversity of SARS-CoV-2

**DOI:** 10.1101/2023.02.10.527437

**Authors:** Ding-Chin Lee, Jui-Hung Tai, Hsin-Fu Lin, Tai-Ling Chao, Yongsen Ruan, Ya-Wen Cheng, Yu-Chi Chou, You-Yu Lin, Sui-Yuan Chang, Pei-Jer Chen, Shiou-Hwei Yeh, Hurng-Yi Wang

## Abstract

Analyses of the genomic diversity of SARS-CoV-2 found that some sites across the genome appear to have mutated independently multiple times with frequency significantly higher than four-fold sites, which can be either due to mutational bias, i.e., elevated mutation rate in some sites of the genome, or selection of the variants due to antagonistic pleiotropy, a condition where mutations increase some components of fitness at a cost to others. To examine how different forces shaped evolution of SARS-CoV-2 in 2020–2021, we analyzed a large set of genome sequences (~ 2 million). Here we show that while evolution of SARS-CoV-2 during the pandemic was largely mutation-driven, a group of nonsynonymous changes is probably maintained by antagonistic pleiotropy. To test this hypothesis, we studied the function of one such mutation, spike M1237I. Spike I1237 increases viral assembly and secretion, but decreases efficiency of transmission *in vitro*. Therefore, while the frequency of spike M1237I may increase within hosts, viruses carrying this mutation would be outcompeted at the population level. We also discuss how the antagonistic pleiotropy might facilitate positive epistasis to promote virus adaptation and reconcile discordant estimates of SARS-CoV-2 transmission bottleneck sizes in previous studies.

## Introduction

Genetic diversity is frequently measured as viral evolvability, making it a useful metric for understanding viral evolution. Genetic diversity is also positively correlated with viral fitness and virulence in emerging viral pathogens. Understanding how viral diversity is generated and maintained is thus crucial in the study of fast-evolving viruses and should contribute to the development of more efficient control and treatment strategies against viral pathogens ^1–4^.

High mutation rates of viral polymerases may underlie much of the adaptive potential of RNA viruses. While *de novo* mutations increase genetic variation, the fate of new genetic variants is largely determined by natural selection. Therefore, the interplay between mutation and selection shapes the genetic diversity. Fitness of a given mutant may be context dependent. For a new mutation generated within a host to be seen in a population, it must adapt to the local environment within a host during infection. In addition, to continually increase its frequency among hosts, the mutant must maintain high transmissibility ^5,6^. However, mutations that increase viral replication rate might affect transmission and/or virulence. The summed effect of selection on all possible effects governs the fate of the mutant ^7^. This antagonistic pleiotropy - where alleles of a gene increase some components of fitness at a cost to others - can contribute to the maintenance of genetic variation ^8–10^.

Since the first report of coronavirus disease 2019 (COVID-19) caused by severe acute respiratory syndrome coronavirus 2 (SARS-CoV-2) in December 2019 in Wuhan, the virus has rapidly sparked an ongoing pandemic ^11–13^. Due to high mutation rate coupled with extraordinary number of infections, SARS-CoV-2 has generated many mutations. Several runs of positive selection have been documented for the origin of D614G, Alpha, Delta, and Omicron variants ^14–16^. The overwhelming number of genome sequences provides an unprecedented opportunity to track evolution in ways unimaginable in the study of any other living organisms. Analyses of the genomic diversity of SARS-CoV-2 found that some sites across the genome appear to have mutated independently multiple times ^17–20^, which can be either due to mutational bias, i.e., elevated mutation rate in some sites of the genome, or selection of the variants due to antagonistic pleiotropy.

Although both of them increase viral diversity, they have different impacts on long-term virus evolution. While elevated mutation rates may mainly generate neutral or deleterious mutations, variations maintained by antagonistic effect have fitness costs and may facilitate epistatic interaction and promote virus adaptation ^21^. Moreover, many aspects of viral dynamics are estimated based on the assumption that the observed variations are selective neutral ^22,23^. If a significant portion of mutations have fitness effects, the accuracy of these estimates would be compromised.

To examine how mutations and different forces of selection, including positive, negative, and antagonistic pleiotropy, shaped evolution of SARS-CoV-2 in 2020–2021, we analyzed a large set of genome sequences (~ 2 million). Our results show that genetic diversity of SARS-CoV-2 during the pandemic is largely mutation driven, as similar mutation spectra were recovered both within and among hosts as well as at synonymous and nonsynonymous sites. While 2/3 of nonsynonymous mutations are deleterious and only a tiny portion (0.3%) of amino acid changes are advantageous, there is a group of nonsynonymous changes with frequency significantly higher than at four-fold degenerate sites that never experienced clonal expansion. These mutations occurred frequently and can reach high frequencies within individuals, indicating they may carry some advantageous effects. However, they might not be highly transmissible, as their frequencies are relatively low in populations, suggesting that antagonistic pleiotropy plays an important role in shaping SARS-CoV-2 genetic variation.

## Results

### SARS-CoV-2 genome sequence evolution is largely mutation-driven

We identified 65,673 SNPs from 28,363 sites out of 29,016 nucleotides in the coding regions of the SARS-CoV-2 genome. Most SNPs are at low frequencies with an average of 4×10^-4^ but a median value of 6.3×10^-6^. The mean frequency among 17,599 synonymous SNPs is 4.6×10^-4^ (median 2.1×10^-5^), while it is 3.8×10^-4^ (median 3.7×10^-6^) among the 48,074 non-synonymous SNPs (Fig. 1A). Significantly lower nonsynonymous than synonymous mutation frequencies indicate that most amino acid changes are deleterious. However, there are certain nonsynonymous mutations that reach significantly higher frequencies than mutations at 4-fold degenerate sites (black dashed line in Fig. 1A), suggesting they are mutational hotspots or under positive or antagonistic selection.

**Figure 1.**
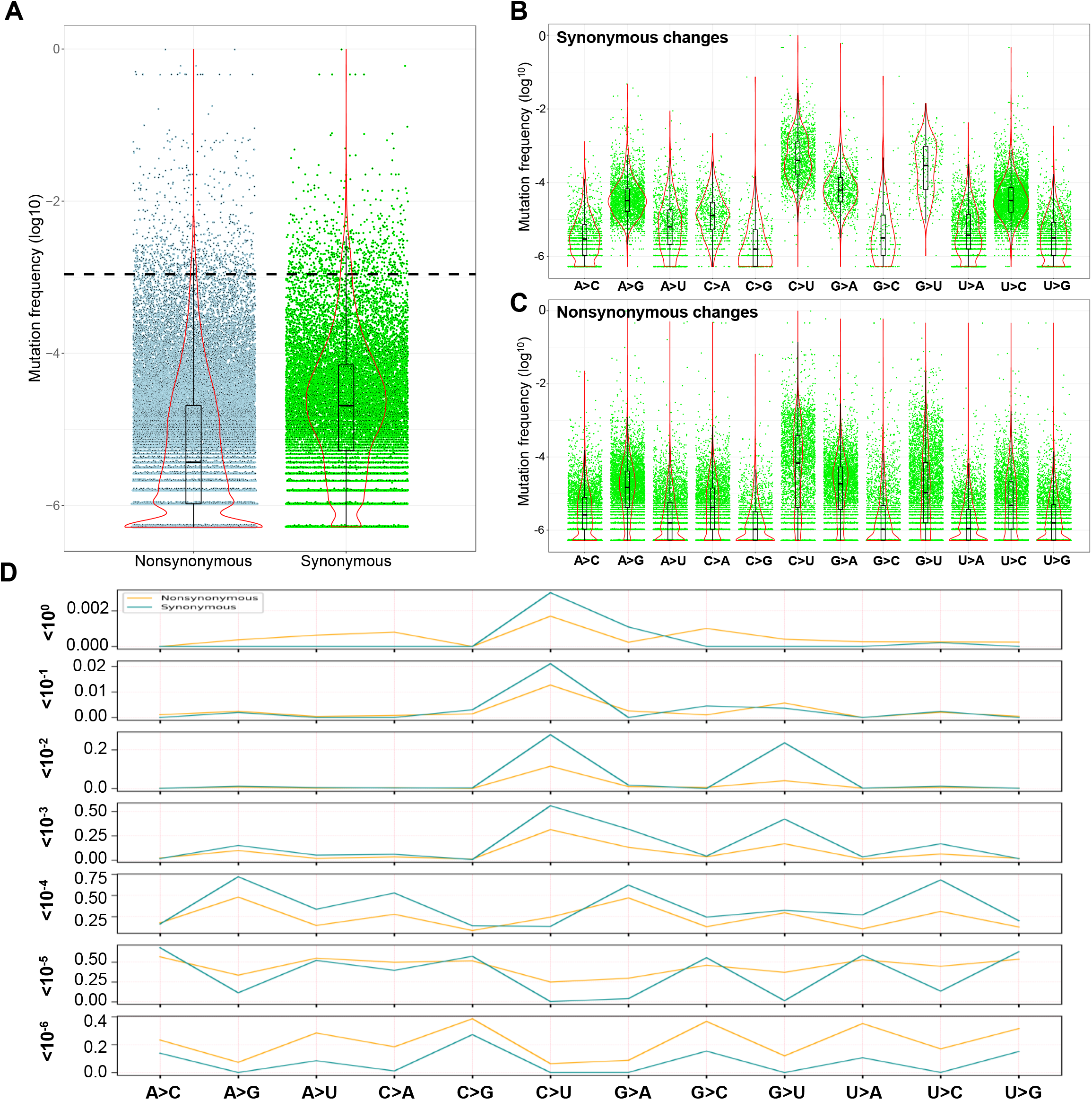
Mutation spectra of SARS-CoV-2. **(A)** Frequency distribution of synonymous and nonsynonymous mutations. Dashed line represents average frequency of fourfold degenerate sites. **(B)** Mutation spectrum of synonymous and **(C)** nonsynonymous mutations. For each mutation type, the violin plot (red) shows probability density of the mutations and the median and the 25th and 75th percentiles of the mutations are shown in boxplots (black). **(D)** Spectra of synonymous and nonsynonymous mutations in frequency bins. The frequencies of synonymous and nonsynonymous mutations are significantly positively correlated in the interval <10^-6^ - <10^-2^ and across all intervals.

Mutational spectra provide insights into the mechanism underlying evolution of SARS-CoV-2. Because our dataset includes most possible mutations at all sites, number of mutations in each category mainly reflect the nucleotide composition of the virus genome and does not directly reflect mutation prevalence. We thus use frequency of nucleotide change as a proxy to estimate the mutation prevalence across types. Cumulative C > U changes reach highest median frequency followed by G > U, G > A, A > G, and U > C changes among synonymous sites (Fig. 1B). Similar patterns are seen for mutations in each month (Fig. S1). These dominant mutation types have been previously described ^19,24,25^. Mutation type reaching highest median frequency among nonsynonymous sites is C > U, followed by G >A, A > G, G > U, and U > C, close to synonymous mutation ranking (Fig. 1C and Fig S2). It was hypothesized that two types of RNA-editing enzyme may contribute to the mutational spectrum. APOBEC, a cytidine deaminase, causing cytosine (C) to uracil (U) transitions, may increase C > U and G >A changes, and ADAR enzymes exchanging adenosine to inosine increase A > G and U > C mutations ^26^. In addition, direct damage of cytosine and guanine bases caused by reactive oxygen species may also contribute to C > U transition and G > U transversion ^18,25^.

When mutations are divided into different intervals according to their frequency, the correlations between synonymous and nonsynonymous mutations are significant in the interval <10^-6^ - <10^-2^ and across all intervals (τ = 0.66; p < 10^-16^; Kendall’s correlation; Fig 1D, Table S1). There are too few mutations with frequency > 10^-2^ to make any inferences. Similar mutational spectra revealed at both synonymous and nonsynonymous mutations suggest that SARS-CoV-2 genome sequence evolution is largely mutation-driven.

Mutational spectra in <10^-6^ and 10^-6^ to 10^-5^ frequency bins are strongly positively correlated for both synonymous and amino acid changing mutations. Interestingly, they are negatively correlated with 10^-5^ to 10^-4^, 10^-4^ to 10^-3^, 10^-3^ to 10^-2^, 10^-2^ to 10^-1^, and 10^-1^ to 10^0^ frequency bins and the latter five frequency bins are also positive correlated with each other (Table S2). Apparently, mutations that are kept under the 10^-5^ frequency exhibit a distinct profile compared to those exceeding this prevalence threshold (Fig 1D).

### Strength of selection against nonsynonymous mutations

The ratio of nonsynonymous (N) to synonymous (S) changes is used to estimate strength of selection. There are roughly 22280.83 nonsynonymous and 6735.17 synonymous sites in the coding regions of the SARS-CoV-2 genome. Thus, with no selection, the N/S ratio of SNPs should be close to 3.31 (22280.83 / 6735.17). A decrease in this ratio can reflect the action of negative selection, while an increase may indicate adaptive changes.

The N/S ratio of SNPs that do not reach 10^-6^ is 11.37, dropping to 4.07 for mutations between 10^-6^ and 10^-5^ (Fig. 2A). The ratios further drop to 1.54 in the 10^-5^ – 10^-4^ bin, 1.40 in the 10^-4^ – 10^-3^ bin, and 1.21 in 10^-3^ – 10^-2^ bin. Therefore, like mutation profiles shown in Fig. 1D, N/S ratios for mutations with frequency < 10^-5^ are very different from those reaching above this threshold. The trend of N/S ratio decreasing is reversed among mutations that exceed 10^-2^ in frequency. We see the same trend of N/S ratio decreasing as mutation frequency goes up (below 1%) when SNPs of different months were analyzed separately (Fig. S3).

**Figure 2.**
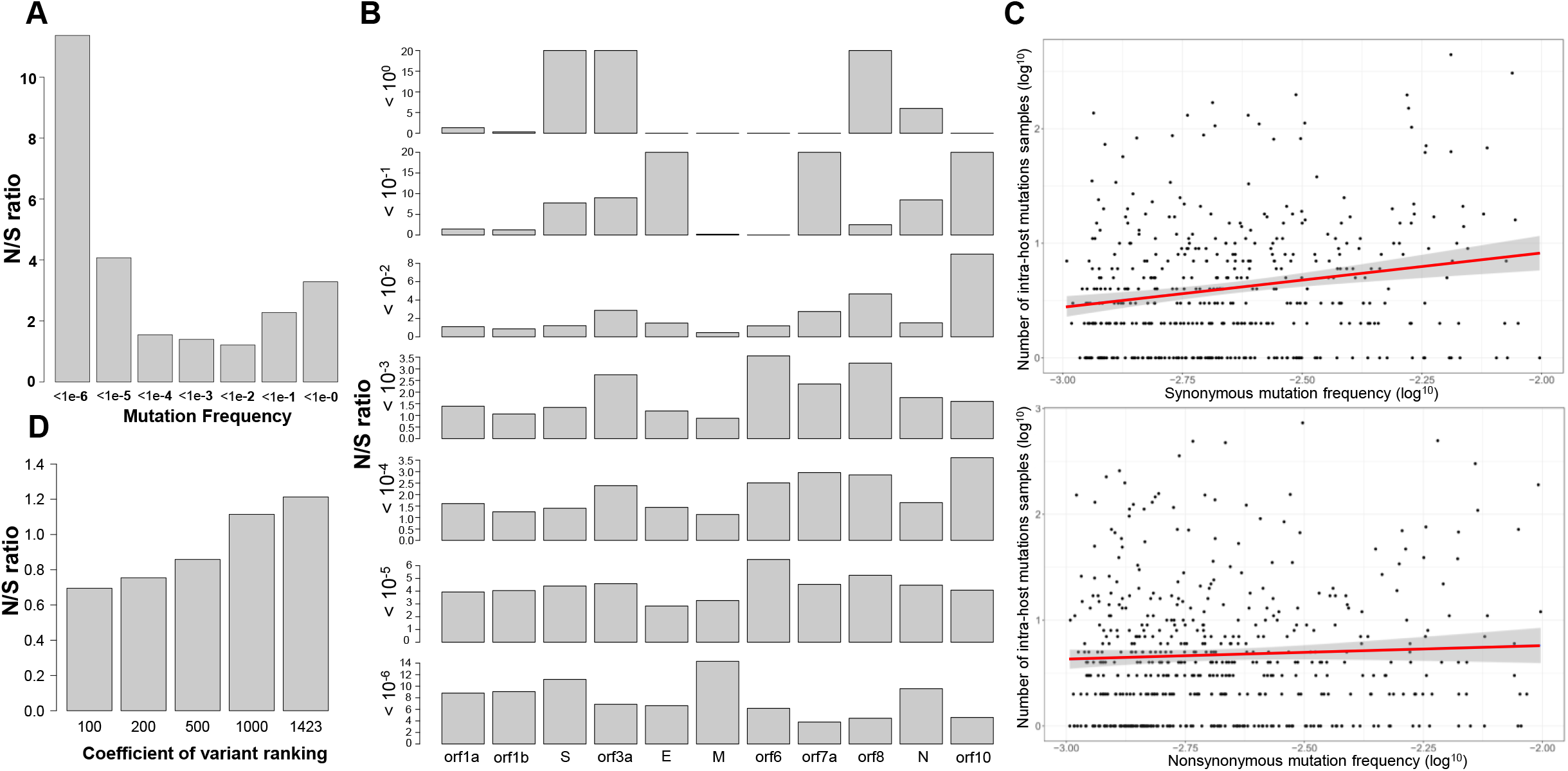
Nonsynonymous (N) and synonymous (S) mutations across frequencies. **(A)** N/S ratios of mutations at different frequencies. **(B)** N/S ratios of mutations at different frequencies across SARS-CoV-2 genome. **(C)** Correlations of synonymous (upper panel, p < 10^-3^, Kendall correlation) and nonsynonymous (lower panel, p = 0.55) mutations with number of intra-host variants (see Materials and Methods). **(D)** N/S ratios of mutations with frequency significantly higher than four-fold degenerate sites ranked by coefficients of variation calculated based on their frequencies across 16 months.

Fig 2A suggests that most amino acid changes with frequency < 10^-5^ are deleterious and only stay at low frequencies, whereas neutral mutations (synonymous changes) may become more prevalent through genetic drift or accumulation of repeated mutations. Indeed, while 65% of nonsynonymous mutations are below 10^-5^, 65% of synonymous mutations are above this cut-off. As a result, 2/3 of the mutations that fail to reach the 10^-5^ frequency exhibit higher N/S ratios than mutations that exceed 10^-5^. In contrast, positive selection drives advantageous mutations to high frequency, increasing the N/S ratio. Consequently, N/S ratios increase again among mutations above 10^-2^.

When ORFs are analyzed separately, most coding regions exhibit high N/S ratios among low frequency mutations (<10^-5^) and the N/S ratios decrease as mutation frequencies increase (Fig. 2B). We see high N/S ratios among envelope and orf7a sites with frequency 10^-2^ – 10^-1^ because there is only one nonsynonymous and no synonymous mutation. The N/S ratios of spike, orf3a, orf8, and nucleocapsid are among highest at low frequency, gradually decreasing as mutation frequencies increase, and rise again among mutations above 10^-2^.

### The role of antagonistic pleiotropy in maintaining high frequency nonsynonymous mutations

We next focus on mutations with frequency higher than the neutral mutation rate. Assuming synonymous mutations are neutral, mutations with frequency significantly higher than the average frequency of 4-fold degenerate sites (1×10^-3^) are considered high frequency (Materials and Methods). This set contains 1630 sites in coding regions. 207 of these sites with frequency > 10^-2^ were removed, as they may be under directional positive selection or genetic hitchhiking as shown in Fig. 2A. The rest are thought to be caused by mutational bias, or, alternatively, maintained by antagonistic pleiotropy.

To test the effect of mutational bias, we analyzed intra-host genomic diversity of SARS-CoV-2 (Materials and Methods). While frequencies of synonymous mutations show strong positive correlation with intra-host variation (p < 10^-3^), frequencies of nonsynonymous mutation do not (p = 0.55) (Fig. 2C). Consequently, while mutational bias may cause some synonymous mutations to reach high frequencies, its effect on nonsynonymous mutations is less obvious.

Antagonistic pleiotropy represents situations where a mutation that has a beneficial effect in one environment has a deleterious effect elsewhere. To test this scenario, we rank the 1423 (1630-207) high frequency mutations according to coefficient of variation (CV) calculated based on their frequencies across 16 months. CV is a measure of dispersion of a frequency distribution. A low CV indicates little frequency fluctuation, as expected under antagonistic interaction or mutational bias. The N/S ratio for the top 100, 200, 500, 1000, and all 1423 sites, are 0.69, 0.75, 0.86, 1.11, and 1.21, respectively (Fig. 2D). High proportion of synonymous mutations with relatively stable frequencies during the course of the pandemic indicates that most of these mutations reflect the underlying mutational bias as previous studies suggested ^18,25^. However, the N/S ratio increases as levels of mutation frequency fluctuation increase, indicating that nonsynonymous mutations in the top-ranking categories are under tremendous selective constrains which in turn suggest that these top-ranking nonsynonymous mutations are maintained by antagonistic pleiotropy.

### S-M1237I increases viral assembly/secretion but decreases infectivity

To test whether high frequency mutations staying at relatively constant frequency over time are maintained by antagonistic pleiotropy, we studied the potential function of mutations in the Spike protein. Two such mutations, M1229I and M1237I, are located at the cytoplasmic tail of the protein, immediately within or proximal to the transmembrane domain (1214 – 1234 a.a.) ^27,28^. This cytoplasmic tail is conserved between SARS-CoV and SARS-CoV-2 (identical in 38 out of 39 a.a., containing nine cysteine residues (Fig. S4)). As documented in above two viruses, mutations within this cytoplasmic tail region might affect both the syncytium formation and viral entry and thus viral infectivity ^29,30^; however, the exact contribution by M1237I in this region remains unclear.

Our previous study showed that the third SARS-CoV-2 outbreak in Taiwan (during April 20 to November 5, 2021) was caused by a single virus lineage (T-III) that carries four genetic fingerprints in the Alpha strain, including spike M1237I (S-M1237I) and three silent changes ^31^. To test the function of S-M1237I, viral titer (determined by plaque forming assay) and viral proteins (determined by immunoblot of lysate from infected cells and from virions in the supernatant) for six SARS-CoV-2 S-M1237-Alpha strains and thirty-two S-I1237-Alpha strains (T-III) isolated at NTUH were compared in Calu-3 cell culture-based virus infectious system (Table S3). The representative immunoblots of intracellular lysates from virus infected Calu-3 cells showed a comparable amount of viral protein in cells infected by these two groups of viruses (Fig. 3A).

**Figure 3.**
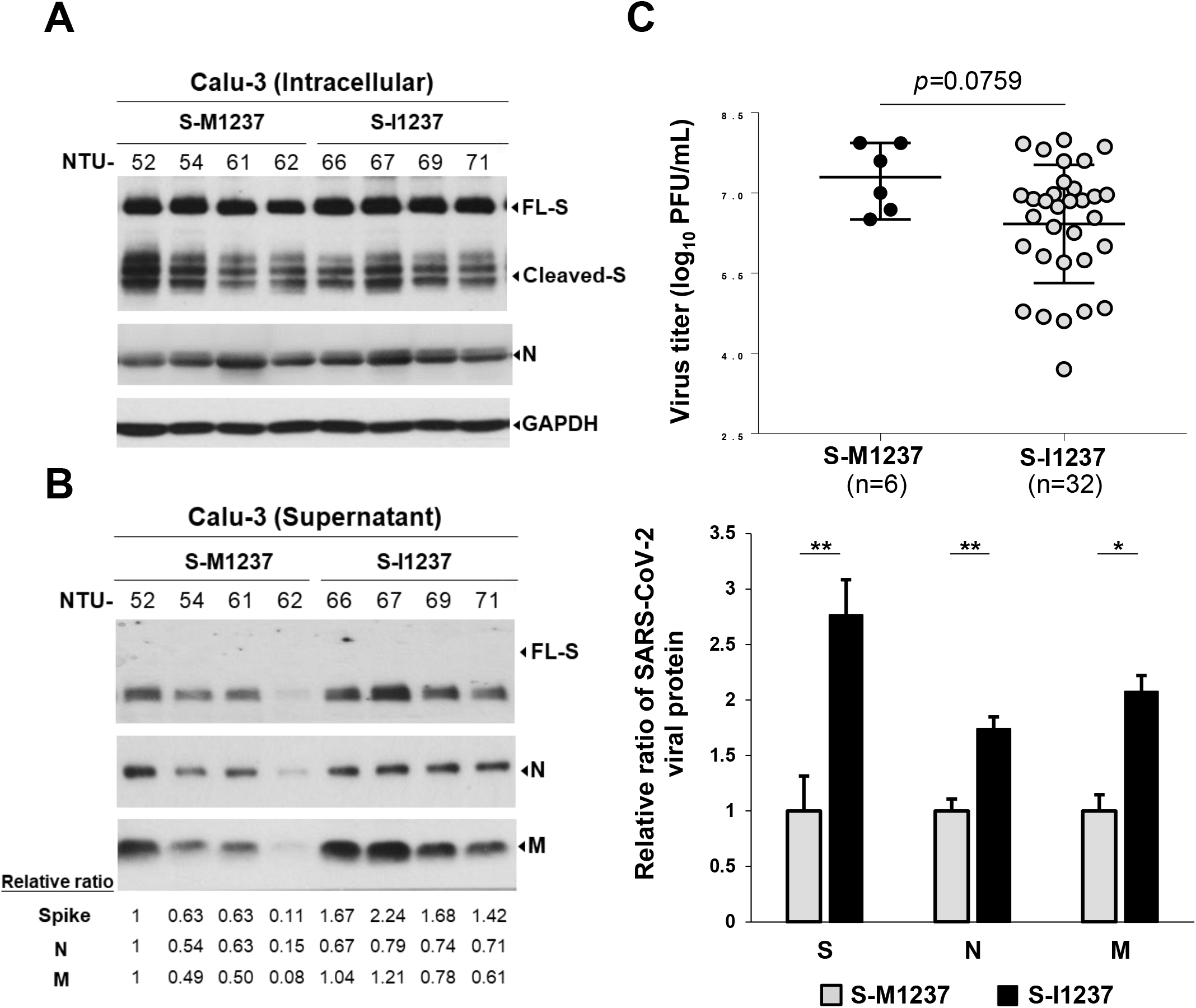
S-M1237I mutation in the SARS-CoV-2 Alpha strains associates with higher viral protein levels but lower plaque forming capability. **(A)(B)** Representative immunoblot results of viral protein expression in Calu-3 cells **(A)** and in supernatants **(B)** after 24hr post infection with SARS-CoV-2 isolates (S-M1237-Alpha and S-I1237-Alpha (T-III) variants, MOI=0.1). The relative ratio of S, N, and M protein in the supernatant was normalized with NTU52 (set as 1). The relative ratios were listed below the immunoblot in left panel and graphically illustrated in right panel of (B). **(C)** Comparison of viral titers (plaque-forming units (PFU)/mL) in the supernatants of Calu-3 cells infected with SARS-CoV-2 isolates ((S-M1237-Alpha and S-I1237-Alpha (T-III) variants, MOI=0.1) at 24 hr post-infection. Data are presented as the mean ± SD (P < 0.01**; P < 0.001***).

A comparison between S-M1237- and S-I1237-Alpha strains showed that the supernatants from the latter had more viral protein but lower plaque forming capability (infectious viral titers) than from the former (Fig. 3B, 3C). This raises the possibility that the assembly/release is more efficient but the infectivity is lower in Alpha strains carrying S-I1237 than in those carrying S-M1237. To address this possibility and validate the critical contribution by the S-M1237I mutation, we compared the viral assembly/release and infection steps individually between two types of reporter virus particles with envelopes containing S-M1237 or S-I1237. The first type is the SARS-CoV-2 spike pseudotyped lentivirus, for analysis of the viral infectivity step (Fig. 4A, upper panel). The second type is the SC2-virus like particles (VLPs), recently developed by Prof. Doudna and Prof. Ott ^32^, for investigating the function of the specific S-M1237I mutation in regulating viral assembly/release and infection steps individually (Fig. 4A, lower panel).

**Figure 4.**
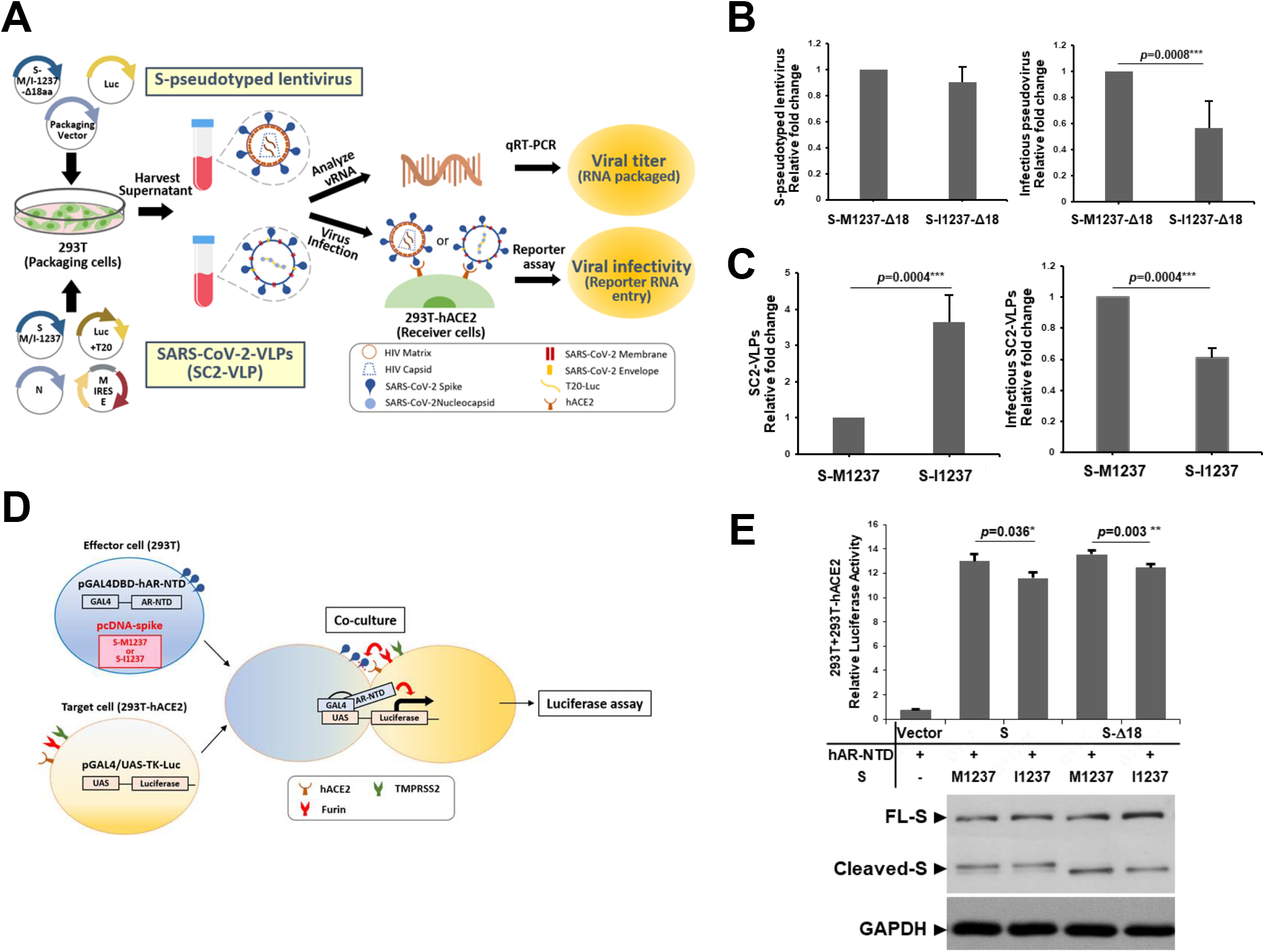
S-M1237I mutation increases viral assembly/secretion but decreases infectivity *in vitro*. **(A)** Schematic illustration of the protocols for generation of S-pseudotyped lentiviruses (upper panel) and SC2-VLPs (lower panel), and for subsequent viral titer and viral infectivity determination. **(B)** Analysis of the viral titer of the lentiviruses pseudotyped with S-B117-M1237-_Δ_18aa or S-B117-I1237-_Δ_18aa produced by 293T cells (left panel) and the viral infectivity (right panel, with S-B117-M1237-_Δ_18aa set as 1). The results were derived from six independent experiments and are shown as the mean ±SD (*P* < 0.05*). **(C)** Analysis of viral titers of the SC2-VLPs containing S-B117-M1237 or S-B117-I1237 produced by 293T cells (left panel) and viral infectivity (right panel, with S-B117-M1237 set as 1). The results were derived from three independent experiments and are shown as the mean ±SD (*P* < 0.05*). **(D)** Schematic illustration of the one-hybrid reporter assay for evaluating fusion activity induced by the SARS-CoV-2 S protein. Effector 293T cells were co-transfected with the expression plasmid for S-B117-M1237 or S-B117-I1237 and the pGAL4DBD-hAR-NTD plasmid. The target 293T-hACE2 cells were transfected with pGAL4/UAS-TK-Luc. At 24 hr post transfection, the effector and target cells were co-cultured for 24 hr and harvested to assay luciferase activity. **(E)** Representative results of the cell-cell fusion activity mediated by S-B117-M1237, S-B117-I1237, S-B117-M1237-**Δ**18aa or S-B117-I1237-**Δ**18aa, which were detected by the luciferase reporter assay of the lysates harvested from co-cultured cells (*P* < 0.05*) (upper panel). The results were derived from three independent experiments and are shown as the mean ± SD (*P* < 0.01**). The expression of the spike protein from 293T cells transfected with the indicated plasmids was analyzed by immunoblotting and GAPDH was included as a loading control (lower panel).

The results from the pseudovirus experiment show a significantly lower viral infectivity of the lentiviruses pseudotyped with S-B117(spike of Alpha strain)-I1237 compared to S-B117-M1237 (Fig. 4B, right panel), though with equivalent viruses released in the supernatant (Fig. 4B, left panel). In this assay, C-terminal 18 amino acids of the ER retention signal in spike were removed (Δ18aa) to optimize pseudotyped lentivirus production ^33^. Since M1237I is the only amino acid different between these two pseudotyped viruses, the results thus directly support the critical function of M1237I in decreasing viral infectivity.

We next generated SC2-VLPs with envelopes containing S-B117-M1237 or S-B117-I1237 and compared their assembly/release and infection efficacy. The expression plasmids for N, M, E and either S-B117-M1237 or S-B117-I1237 were co-transfected with a packaging signal containing luciferase-encoding mRNA, Luc-T20, into the packaging 293T cells. Quantification of this Luc-T20 RNA packaged and released to the supernatant by qRT-PCR shows that more mRNA-containing SC2-VLPs were released from the cells expressing S-B117-I1237 than from S-B117-M1237 producing cells (Fig. 4C, left panel). Therefore, the S-M1237I mutation might contribute to the increase of SARS-CoV-2 assembly and release.

The mRNA-containing VLPs in the supernatant were then harvested for infection of the receiver ACE2-expressing 293T cells. We measured luciferase activity from the cells infected with the same amount of Luc-T20-containing SC2-VLPs (MOI=0.05) to estimate infectivity rates. S-B117-I1237 containing SC2-VLPs show significantly less luciferase than S-B117-M1237 SC2-VLPs, indicating lower infectivity (Fig. 4C, right panel). These results altogether suggest that S-M1237I contributes to an increase of assembly/release but a decrease of infectivity in T-III Alpha strain viruses.

To further investigate the underlying mechanism for this lower infectivity, we examined viral-cell membrane fusion mediated by spike-ACE2 interaction using the syncytium formation assay (details schematically illustrated in Fig. 4D). We found significantly lower syncytium formation activity mediated by S-B117-I1237 than S-B117-M1237 in both full-length spike and Δ18aa spike proteins (Fig. 4E). Since viral-cell membrane fusion is the critical step determining viral infectivity, these results suggest that S-M1237I contributes to the lower infectivity of T-III strains by decreasing viral-cell membrane fusion activity.

## Discussion

By analyzing 1,929,395 SARS-CoV-2 genomes, we identified 17,599 synonymous and 48,074 nonsynonymous mutations and used them to dissect forces that drive SARS-CoV-2 evolution during the pandemic. We find similar variation spectra within and between hosts. Several mechanisms, including APOBEC enzyme increasing C > U and G >A changes, ADAR enzymes which may increase A>G and U > C mutations, and reactive oxygen species that contribute to C > U transition and G > U transversion, may cause biased mutational spectra within hosts ^24,26,34–37^. These five dominant mutational patterns appear to comprise mutation profiles within populations for both nonsynonymous and synonymous changes (Fig. 1B and 1C). Moreover, mutational spectra for two types of changes at different frequencies are all highly correlated (Fig. 1D, Table S1), demonstrating that evolution of SARS-CoV-2 during the pandemic is largely mutation-driven.

We find genetic variation patterns in SARS-CoV-2 are dominated by negative selection ^38^, as the median frequency of synonymous mutations is five time higher than nonsynonymous mutations (Fig. 1A). In addition, 2/3 of nonsynonymous mutations have their frequency < 10^-5^, whereas only 1/3 of synonymous mutations are at that level. While neutral mutations (synonymous changes) may increase their frequency by genetic drift or accumulation of repeated mutations, deleterious mutations can only stay at low frequency as they are subjected to negative selection. As a result, N/S ratios for mutations with frequency < 10^-5^ are highest, indicating that nonsynonymous mutations at these frequencies are largely deleterious. In contrast, high N/S ratios are also seen in the > 10^-2^ bin. As positive selection increases the frequencies of advantageous mutants, the overrepresentation of nonsynonymous mutations in this high-prevalence bin indicates that they are adaptive. Consequently, we estimate that 2/3 of nonsynonymous mutations are deleterious and only 0.3% (at frequencies above 10^-2^) are advantageous.

The above result may suggest that most remaining 1/3 of nonsynonymous mutations are neutral. However, there is a group of mutations with frequencies > 1×10^-3^ that is significantly higher than 4-fold degenerate sites. It is possible that mutational bias increases mutation rates at these sites. Synonymous site mutation frequencies at the population level are significantly positively correlated with within host prevalence, suggesting high mutation load at these sites (Fig. 2C). However, such correlation was not seen for nonsynonymous sites. We thus hypothesize that high mutation frequency of these nonsynonymous mutations is maintained by antagonistic pleiotropy, representing situations in which a mutation that has a beneficial effect in one environment has a deleterious effect elsewhere. This is expected for organisms, such as viruses, that constantly shift between different environments, such as spreading among hosts and adapting within hosts. Besides, their mutation frequency may be relatively constant over time, as the viruses are shifting between and within hosts.

To test the above hypothesis, we studied the function of the spike M1237I mutation. We found that SARS-CoV-2 carrying the spike I1237 variant secretes more viral particles than its counterparts carrying M1237. However, this comes at a cost to infectivity. Demographically, M1237I is found in ~0.13% of all SARS-CoV-2 genomes and is therefore not a rare mutation and can be commonly found in different genetic backgrounds (Table S4), indicating that it was repeatedly and frequently created. This can be explained by the dual demand (i.e., adaptation within hosts and colonization between hosts) of the viral life cycle ^6^. Within hosts, SARS-CoV-2 with spike I1237 may rapidly increase its frequency, because it secretes more viral particles than its peers. As a result, we frequently observe the rise of spike M1237I in some individuals. However, because spike I1237 may decrease virus infectivity, SARS-CoV-2 carrying this mutation will be surpassed by those carrying M1237 at the population level. Therefore, this mutation is usually quickly eliminated. If this explanation is correct, we would expect viruses carrying spike I1237 to continuously spread providing that there is no other strain competing with it. Indeed, during the third outbreak of SARS-CoV-2 which resulted in 13,795 cases in Taiwan in 2021, all viruses sequenced carried spike I1237, indicating this mutation is capable of transmission. As there was no alternative strain in Taiwan during the third outbreak, the virus strain was thus continuously transmitted within the community ^31^.

Our study has implications for our understanding of virus evolution. First, in order to continually spread among hosts, the virus must maintain high transmissibility. On the other hand, the virus must adapt to the local environment within a host during the course of infection, even at the cost of lowered transmissibility. Some mutations might be advantageous at transmission, whereas others might be good at replication within hosts. However, it is difficult for a single mutation to be advantageous at transmission, assembly, and replication. Accordingly, a single mutation is difficult thriving. In order to realize the fitness effect, a group of mutations must be in strong epistatic interaction. For example, it has been proposed that while many Omicron BA.1 Spike mutations are likely to be maladaptive when present on their own, collectively they might interact with each other to reduce the deleterious effect, i.e., antagonistic epistasis ^21^, or even compensate for one another’s deficits to build a better virus, i.e., positive epistasis ^16^. Antagonistic pleiotropy which maintains diversity within population may facilitate epistatic interaction among different mutations. If this implication is correct, we would expect many of the Omicron mutations coincide with high frequency mutations (10^-3^ ~ 10^-2^) identified by this study. Among 46 amino acid changes in Omicron BA. 2, 12 of them have frequency > 10^-2^ in our dataset, suggesting that they are under positive selection. Eleven out of the rest 34 changes are in the frequency bin of 10^-3^ – 10^-2^ (odds ratio = 28.9, p < 10^-11^, Fisher exact test; Table S5), demonstrating that many of high frequency mutations are probably under antagonistic pleiotropy.

Second, our results may explain the discordant estimates of SARS-CoV-2 transmission bottleneck sizes in previous studies, which ranged from 1-10 virions to 100 - 1,000 virions ^20,25,39–41^. That is because the current method for transmission bottleneck size estimation considers all mutations within hosts are neutral ^22,42^. The antagonistic pleiotropy implies that there are two layers of selection, i.e., within and between hosts, during virus evolution. The probability for a mutation to be transmitted is determined not only by the size of transmission bottleneck as neutral model predicted, but also by its effect at different stages as suggested by our study of S-M1237I. Consequently, mutations with antagonistic pleiotropy would bias the estimation of bottleneck size. As accurate quantification of transmission bottleneck size will help in forecasting the rate of adaptation for rapidly evolving pathogen such as SARS-CoV-2 ^22,23^, it is therefore necessary to consider effect of antagonistic pleiotropy during viral transmission in the future.

## Methods

### SARS-CoV-2 genome collection and analysis

The data collection and preprocessing are as previously described ^43^. In brief, we downloaded SARS-CoV-2 genomes from the GISAID database (https://www.gisaid.org/) as of July 5, 2021. We aligned these 1,929,395 genome sequences to the reference sequence (EPI_ISL_402125) using MAFFT^44^ (--auto --keeplength). We used snp-sites (-v; Page et al. 2016) to identify single nucleotide polymorphisms (SNPs) and BCFtools ^45^ (merge–force-samples -O v) to merge the vcf files. We identified 65,673 SNPs in coding regions. The SNPs were classified into 12 mutation types as shown in Fig. 1.

To define high frequency mutations, we first calculated average mutation frequency of four-fold degenerate sites (7.4×10^-4^). With 4236 four-fold degenerate sites and standard deviation of 1.26×10^-2^, the 95% confidence interval for four-fold degenerate site frequency is 3.6×10^-4^ to 1.1 ×10^-3^. We consider mutations with frequency > 10^-3^ as high frequency for convenience.

For intra-host variation of SARS-CoV-2, publicly available high-throughput sequencing data sets were downloaded from the NCBI Sequence Read Archive to assess intra-host genetic variation. Data set IDs are listed in Table S6. Sequence reads were mapped to the Wuhan-Hu-1 reference genome (EPI_ISL_402125) using the Burrows-Wheeler Aligner (BWA) ^45^. Nucleotide frequencies at each position were extracted from files generated by *bcftools* ^46^. To reduce the number of variants due to sequencing errors, only sequencing depths greater than 20 reads and mapping quality > 20 were considered. All further calculations were performed using R and in-house Python scripts.

### SARS-CoV-2 viruses

Thirty-eight virus isolates were obtained from the sputum specimens of SARS-CoV-2-infected patients, propagated in Vero E6 cells in Dulbecco’s modified Eagle’s medium (DMEM) supplemented with 2 μg/mL tosylsulfonyl phenylalanyl chloromethyl ketone-trypsin. Virus isolates used in the current study have been deposited in the GISAID platform (available at https://www.gisaid.org/). Accession numbers are listed in Table S3.

### Plaque forming assay

The plaque assay was performed following the protocol described previously ^47^. In brief, Vero E6 cells (2×10^5^ cells/well) were seeded in triplicate in 24-well tissue culture dishes in DMEM supplemented with 10% fetal bovine serum (FBS) and antibiotics. After 24 hrs of infection, virus-containing culture supernatant was added to the cell monolayer for 1 hr at 37°C, which was washed with PBS and maintained with the medium containing 1% methylcellulose. After incubation for 5 days, cells were fixed with 10% formaldehyde overnight and stained with 0.7% crystal violet for plaque counting. Plaque forming activity is estimated from three independent experiments.

### Plasmid constructs

The humanized SARS-CoV-2 spike expression plasmid in pcDNA3.0-HA vector, SCoV2-S, and the mutated SCoV2-S-D614G was constructed as described previously ^48^. To construct the expression plasmids for the spike of Alpha variant (B.1.1.7), pcDNA3.1(+)-SCoV2-S(B.1.1.7), we purchased the synthetic DNA fragment encoding spike of the Alpha variant virus (B.1.1.7) from Integrated DNA Technologies (IDT) for cloning into the kpnI and EcoRI restriction enzyme sites of pcDNA3.1(+) expression vector (Thermofisher). The construct contains the representative mutations of the Alpha variant virus, with 69-70 del, Y144 del, N501Y, A570D, D614G, P681H, T716I, S982A and D1118H. In addition, a pcDNA3.1(+)-SCoV2-S(B.1.1.7)-Δd18aa construct, where the C-terminal 18 amino acids of the ER retention signal in spike were removed to optimize pseudotyped lentivirus production ^33^. The mutant M1237I spike related constructs were generated using the QuikChange II site-directed mutagenesis kit (Agilent) with the primer set of S-M1237I-F: 5’-ACAGCAGGATGTTATACAGCACAGCATGATGGTCAC-3’ and S-M1237I-R: 5’-GTGACCATCATGCTGTGCTGTATAACATCCTGCTGT-3’. The plasmids expressing SARS-CoV-2 membrane or envelope protein (pLVX-EF1alpha-SARS-CoV-2-M and pLVX-EF1alpha-SARS-CoV-2-E) with 2x streptavidin tag at the C-terminus were kindly provided by Dr. Chia-Wei Li from the Institute of Biomedical Sciences Center, Academia Sinica, Taiwan.

### Cell culture experiments

The 293T cells stably expressing human ACE2 (293T-hACE2) were kindly provided by Dr. Mi-Hua Tao (Institute of Biomedical Sciences Center, Academia Sinica, Taiwan). The 293T, 293T-hACE2 and Calu-3 cells were maintained in DMEM medium contained 10-15% FBS as described ^49^. The plasmid was transfected into 293T cells with Lipofectamine™ 2000 (Thermo Fisher Scientific) and the cells were harvested 24 hr post transfection for subsequent analysis.

### Cell-cell fusion assay

A quantitative GAL4-based mammalian luciferase reporter assay was established to assess cellcell fusion activity as described previously, containing a reporter construct (pGAL4/UAS-TK-Luc) ^48^ and a transcriptional activator construct (pGAL4DBD-hAR-NTD). To detect cell-cell fusion activity mediated by WT-S and M1237I-S protein, 293T-hACE2 cells transfected with pGAL4/UAS-TK-Luc were prepared as target cells; 293T cells expressing the either WT-S or M1237I-S and pGAL4DBD-hAR-NTD were prepared as effector cells. After 24 hr transfection, 293T cells expressing the GAL4DBD-hAR-NTD protein and S protein were suspended by trypsin, and seeded on 293T-hACE2 cells expressed the GAL4/UAS-TK-Luc protein (293T: 293T-hACE2 ratio=1:3). The cells were co-cultured for 20 hrs and harvested to measure the fusion activity by detecting the luciferase assay following the manufacture’s instruction (Promega).

### Production and purification of SARS-CoV-2 spike pseudotyped lentivirus

The pseudotyped lentivirus carrying various SARS-CoV-2 spike proteins was generated by transiently transfecting 293T cells with pCMV-DR8.91, pLAS2w.Fluc.Ppuro and pcDNA3.1-SCoV2-S(B.1.1.7) related spike expression constructs using TransITR-LT1 transfection reagent (Mirus). Culture media were refreshed at 16 hr and harvested at 48 hr and 72hr post-transfection. Cell debris was removed by centrifugation at 4,000 xg for 10 min; the supernatant was passed through 0.45-mm syringe filter (Pall Corporation) and the pseudotyped lentivirus was aliquoted and stored at −80°C.

### Estimation of lentiviral titer using the alarmaBlue assay

The transduction unit (TU) of SARS-CoV-2 peudotyped lentivirus was estimated by using a cell viability assay in response to limited lentivirus dilution. In brief, 293T-hACE2 cells (stably expressing human ACE2) were plated on 96-well plates one day before lentivirus transduction. To determine the titer of the pseudotyped lentivirus, different amount of lentivirus was added into the culture medium containing polybrene (final concentration 8 mg/ml). Spin infection was carried out at 1,100 xg in 96-well plate for 30 minutes. After incubating cells at 37°C for 16 hr, the culture medium containing virus and polybrene were removed and replaced with complete DMEM containing 2.5 μg/ml puromycin. After treating with puromycin for 48 hrs, the culture media were removed and cell viability was estimated using 10% AlarmaBlue reagents according to manufacturer’s instruction (ThermoFisher). The survival rate of uninfected cells (without puromycin treatment) was set as 100%. Virus titers (transduction units) were determined by plotting cell survival against diluted viral dose.

### SARS-CoV-2-virus like particle (SC2-VLP) production and infection

SC2-VLPs containing luciferase (Luc)-encoding transcripts enveloped with S-I1237 or S-M1237 were prepared as previously described ^32^, with minor modifications. In brief, the Luc-T20 expressing construct, encoding the Luc mRNA with the SARS-CoV-2 packaging signal in 3’-UTR (Addgene), were co-transfected with plasmids expressing the SARS-CoV-2 nucleocapsid (nCoV-2-N), membrane and envelope (CoV2-M-IRES-E, Addgene), and spike (nCoV-2-B117 or B117/M1237I) into the packaging 293T cells (with molar ratio for Luc-T20: N: M/E: S as 3: 1: 1: 1) with Lipofectamine^TM^ 2000 (Thermo Fisher Scientific). At 24 and 48 hours post-transfection, culture media were collected and filtered through 0.45 μm filters, which were processed to determine viral titer and infectivity. For viral titer determination, the supernatant was first treated with 6000 U micrococcal nuclease (NEB) before viral RNA extraction with MagNA Pure (Roche Diagnostics), which were reverse transcribed using SuperScript III Reverse Transcriptase System (Thermo Fisher Scientific). Quantitative PCR was performed using FastStart DNA SYBR Green on LightCycler 1.5 (Roche Diagnostics), with primer set of 5’-AGACAGTGGTTGCCTACGGG-3’ and 5’-ATGCGAAGTGTCCCATGAGC-3’. For infectivity determination, the supernatant containing equal amounts of Luc-VLP (MOI=0.05) was processed to infect human ACE2-expressing 293T cells. 24 hours later, the cells were lysed using a passive lysis buffer (Promega) and equal amounts of lysates were used for luciferase reporter assay following the manufacturer’s instructions (Promega).

### Western blot analysis

Western blotting was performed using sodium dodecyl sulfate polyacrylamide gel electrophoresis (SDS-PAGE) and Western Lightning Plus-ECL (PerkinElmer) as previously described ^50^. Antibodies used for immunoblotting are as follows: rabbit anti-SCoV/SARS-CoV-2 nucleocapsid (generated by our laboratory), mouse anti-SARS-CoV/SARS-CoV-2 (COVID-19) spike [1A9] (Genetex, GTX632604), rabbit anti-SARS-CoV-2 membrane (Novus Biologicals, NBP3-05698), rabbit anti-SARS-CoV-2 envelope (Cell signaling, #74698), rabbit anti-GAPDH (Genetex, GTX100118), horseradish peroxidase-conjugated mouse IgG (Genetex, GTX213111-01) and rabbit IgG (Genetex, GTX213110-01). Quantitative comparisons of viral protein amount on the blots were made using VisionWorks Life Science Image Analysis software (UVP, Upland, CA USA).

### Statistical analysis

The plaque forming units quantified by plaque assays in triplicate are shown as the mean ± SD. Results from the cell-cell fusion reporter assay are shown as data representative of three independent experiments and presented as mean ± SD. Differences in data from the virus titer and fusion reporter assay between each indicated paired groups were evaluated by Student’s t-test. A P value <= 0.05 was considered statistically significant (*, *P* < 0.05; **, *P* < 0.01; ***, *P* < 0.001).

## Acknowledgement

This study was supported by grants from the Ministry of Science and Technology (MOST), Taiwan (111-2321-B-002-017, 111-2634-F-002-017, 109-2311-B-002-023-MY3) and the “Center of Precision Medicine” from The Featured Areas Research Center Program within the framework of the Higher Education Sprout Project by the Ministry of Education (MOE) in Taiwan.

## Author contributions

JHT, HFL, and YYL analyzed the data. DCL, TLC, YWC, and YCC conducted the experiments. SYC, PJC, SHY, and HYW design the study, obtained funding, and wrote the manuscript.

## Reference

1 Rahimi, A., Mirzazadeh, A. & Tavakolpour, S. Genetics and genomics of SARS-CoV-2: A review of the literature with the special focus on genetic diversity and SARS-CoV-2 genome detection. Genomics 113, 1221–1232, doi:https://doi.org/10.1016/j.ygeno.2020.09.059 (2021).

2 Borderia, A. V., Stapleford, K. A. & Vignuzzi, M. RNA virus population diversity: implications for inter-species transmission. Current Opinion in Virology 1, 643–648, doi:10.1016/j.coviro.2011.09.012 (2011).

3 Duffy, S. Why are RNA virus mutation rates so damn high? PLOS Biology 16, e3000003, doi:10.1371/journal.pbio.3000003 (2018).

4 Mattenberger, F., Vila-Nistal, M. & Geller, R. Increased RNA virus population diversity improves adaptability. Scientific Reports 11, 6824, doi:10.1038/s41598-021-86375-z (2021).

5 Pybus, O. G. & Rambaut, A. Evolutionary analysis of the dynamics of viral infectious disease. Nature Reviews Genetics 10, 540–550, doi:10.1038/nrg2583 (2009).

6 Lin, Y. Y. et al. New Insights into the Evolutionary Rate of Hepatitis B Virus at Different Biological Scales. Journal of Virology 89, 3512–3522, doi:10.1128/Jvi.03131-14 (2015).

7 Day, T., Gandon, S., Lion, S. & Otto, S. P. On the evolutionary epidemiology of SARS-CoV-2. Current Biology 30, R849–R857, doi:10.1016/j.cub.2020.06.031 (2020).

8 Presloid, J. B., Ebendick-Corpus, B. E., Zarate, S. & Novella, I. S. Antagonistic pleiotropy involving promoter sequences in a virus. Journal of Molecular Biology 382, 342–352, doi:10.1016/j.jmb.2008.06.080 (2008).

9 Hedrick, P. W. Antagonistic pleiotropy and genetic polymorphism: a perspective. Heredity 82, 126–133, doi:10.1038/sj.hdy.6884400 (1999).

10 Connallon, T. & Chenoweth, S. F. Dominance reversals and the maintenance of genetic variation for fitness. PLOS Biology 17, e3000118, doi:10.1371/journal.pbio.3000118 (2019).

11 Chaw, S.-M. et al. The origin and underlying driving forces of the SARS-CoV-2 outbreak. Journal of Biomedical Science 27, 73, doi:10.1186/s12929-020-00665-8 (2020).

12 Tang, X. et al. On the origin and continuing evolution of SARS-CoV-2. National Science Review 7, 1012–1023, doi:10.1093/nsr/nwaa036 (2020).

13 Wu, F. et al. A new coronavirus associated with human respiratory disease in China. Nature 579, 265–269, doi:10.1038/s41586-020-2008-3 (2020).

14 Tegally, H. et al. Detection of a SARS-CoV-2 variant of concern in South Africa. Nature 592, 438–443, doi:10.1038/s41586-021-03402-9 (2021).

15 Korber, B. et al. Tracking Changes in SARS-CoV-2 Spike: Evidence that D614G Increases Infectivity of the COVID-19 Virus. Cell 182, 812–827.e819, doi:https://doi.org/10.1016/i.cell.2020.06.043 (2020).

16 Martin, D. P. et al. Selection Analysis Identifies Clusters of Unusual Mutational Changes in Omicron Lineage BA.1 That Likely Impact Spike Function. Molecular Biology and Evolution 39, msac061, doi:10.1093/molbev/msac061 (2022).

17 van Dorp, L. et al. Emergence of genomic diversity and recurrent mutations in SARS-CoV-2. Infection, Genetics and Evolution 83, 104351, doi:https://doi.org/10.1016/j.meegid.2020.104351 (2020).

18 Tonkin-Hill, G. et al. Patterns of within-host genetic diversity in SARS-CoV-2. Elife 10, doi:10.7554/eLife.66857 (2021).

19 Li, J. et al. Two-step fitness selection for intra-host variations in SARS-CoV-2. Cell Reports 38, 110205, doi:10.1016/j.celrep.2021.110205 (2022).

20 Lythgoe, K. A. et al. SARS-CoV-2 within-host diversity and transmission. Science 372, eabg0821, doi:10.1126/science.abg0821 (2021).

21 Desai, M. M., Weissman, D. & Feldman, M. W. Evolution Can Favor Antagonistic Epistasis. Genetics 177, 1001–1010, doi:10.1534/genetics.107.075812 (2007).

22 Leonard, A. S., Weissman, D. B., Greenbaum, B., Ghedin, E. & Koelle, K. Transmission Bottleneck Size Estimation from Pathogen Deep-Sequencing Data, with an Application to Human Influenza A Virus. Journal of Virology 91, e00171–00117, doi:doi:10.1128/JVI.00171-17 (2017).

23 McCrone, J. T. & Lauring, A. S. Genetic bottlenecks in intraspecies virus transmission. Current Opinion in Virology 28, 20–25, doi:https://doi.org/10.1016/j.coviro.2017.10.008 (2018).

24 Simmonds, P. & Schwemmle, M. Rampant C->U Hypermutation in the Genomes of SARS-CoV-2 and Other Coronaviruses: Causes and Consequences for Their Short- and Long-Term Evolutionary Trajectories. mSphere 5, e00408–00420, doi:doi:10.1128/mSphere.00408-20 (2020).

25 Graudenzi, A., Maspero, D., Angaroni, F., Piazza, R. & Ramazzotti, D. Mutational signatures and heterogeneous host response revealed via large-scale characterization of SARS-CoV-2 genomic diversity. iScience 24, 102116, doi:10.1016/j.isci.2021.102116 (2021).

26 Di Giorgio, S., Martignano, F., Torcia, M. G., Mattiuz, G. & Conticello, S. G. Evidence for host-dependent RNA editing in the transcriptome of SARS-CoV-2. Science Advances 6, eabb5813, doi:10.1126/sciadv.abb5813 (2020).

27 Buonvino, S. & Melino, S. New Consensus pattern in Spike CoV-2: potential implications in coagulation process and cell–cell fusion. Cell Death Discovery 6, 134, doi:10.1038/s41420-020-00372-1 (2020).

28 Sanders, D. W. et al. SARS-CoV-2 requires cholesterol for viral entry and pathological syncytia formation. eLife 10, e65962, doi:10.7554/eLife.65962 (2021).

29 Puthenveetil, R. et al. S-acylation of SARS-CoV-2 spike protein: Mechanistic dissection, in vitro reconstitution and role in viral infectivity. Journal of Biological Chemistry 297, ARTN 101112, doi:10.1016/j.jbc.2021.101112 (2021).

30 Li, D. Q., Liu, Y. H., Lu, Y., Gao, S. & Zhang, L. L. Palmitoylation of SARS-CoV-2 S protein is critical for S-mediated syncytia formation and virus entry. Journal of Medical Virology 94, 342–348, doi:10.1002/jmv.27339 (2022).

31 Tai, J.-H. et al. Spatial and temporal origin of the third SARS-COV-2 Outbreak in Taiwan. bioRxiv, 2022.2007.2004.498645, doi:10.1101/2022.07.04.498645 (2022).

32 Syed, A. M. et al. Rapid assessment of SARS-CoV-2-evolved variants using virus-like particles. Science 374, 1626–1632, doi:doi:10.1126/science.abl6184 (2021).

33 Xiong, H.-L. et al. Robust neutralization assay based on SARS-CoV-2 S-protein-bearing vesicular stomatitis virus (VSV) pseudovirus and ACE2-overexpressing BHK21 cells. Emerging Microbes & Infections 9, 2105–2113, doi:10.1080/22221751.2020.1815589 (2020).

34 Krokan, H. E., Drabløs, F. & Slupphaug, G. Uracil in DNA – occurrence, consequences and repair. Oncogene 21, 8935–8948, doi:10.1038/sj.onc.1205996 (2002).

35 Helleday, T., Eshtad, S. & Nik-Zainal, S. Mechanisms underlying mutational signatures in human cancers. Nature Reviews Genetics 15, 585–598, doi:10.1038/nrg3729 (2014).

36 Kustin, T. & Stern, A. Biased Mutation and Selection in RNA Viruses. Molecular Biology and Evolution 38, 575–588, doi:10.1093/molbev/msaa247 (2020).

37 Takata, M. A. et al. CG dinucleotide suppression enables antiviral defence targeting non-self RNA. Nature 550, 124–127, doi:10.1038/nature24039 (2017).

38 Morales, A. C. et al. Causes and Consequences of Purifying Selection on SARS-CoV-2. Genome Biology and Evolution 13, doi:10.1093/gbe/evab196 (2021).

39 Wang, D. et al. Population Bottlenecks and Intra-host Evolution During Human-to-Human Transmission of SARS-CoV-2. Frontiers in Medicine 8, doi:10.3389/fmed.2021.585358 (2021).

40 Popa, A. et al. Genomic epidemiology of superspreading events in Austria reveals mutational dynamics and transmission properties of SARS-CoV-2. Science Translational Medicine 12, eabe2555, doi:doi:10.1126/scitranslmed.abe2555 (2020).

41 Hannon, W. W. et al. Narrow transmission bottlenecks and limited within-host viral diversity during a SARS-CoV-2 outbreak on a fishing boat. Virus Evolution 8, doi:10.1093/ve/veac052 (2022).

42 Braun, K. M. et al. Transmission of SARS-CoV-2 in domestic cats imposes a narrow bottleneck. PLOS Pathogens 17, e1009373, doi:10.1371/journal.ppat.1009373 (2021).

43 Ruan, Y. et al. The Runaway Evolution of SARS-CoV-2 Leading to the Highly Evolved Delta Strain. Molecular Biology and Evolution 39, doi:10.1093/molbev/msac046 (2022).

44 Katoh, K. & Standley, D. M. MAFFT Multiple Sequence Alignment Software Version 7: Improvements in Performance and Usability. Molecular Biology and Evolution 30, 772–780, doi:10.1093/molbev/mst010 (2013).

45 Li, H. & Durbin, R. Fast and accurate short read alignment with Burrows-Wheeler transform. Bioinformatics 25, 1754–1760, doi:10.1093/bioinformatics/btp324 (2009).

46 Danecek, P. et al. Twelve years of SAMtools and BCFtools. GigaScience 10, doi:10.1093/gigascience/giab008 (2021).

47 Su, C. T. et al. Anti-HSV activity of digitoxin and its possible mechanisms. Antiviral Res 79, 62–70, doi:10.1016/j.antiviral.2008.01.156 (2008).

48 Cheng, Y.-W. et al. D614G Substitution of SARS-CoV-2 Spike Protein Increases Syncytium Formation and Virus Titer via Enhanced Furin-Mediated Spike Cleavage. mBio 12, e00587–00521, doi:10.1128/mBio.00587-21 (2021).

49 Cheng, Y.-W. et al. Furin Inhibitors Block SARS-CoV-2 Spike Protein Cleavage to Suppress Virus Production and Cytopathic Effects. Cell Reports 33, 108254, doi:https://doi.org/10.1016/j.celrep.2020.108254 (2020).

50 Wu, C. H. et al. Glycogen synthase kinase-3 regulates the phosphorylation of severe acute respiratory syndrome coronavirus nucleocapsid protein and viral replication. Journal of Biological Chemistry 284, 5229–5239, doi:10.1074/jbc.M805747200 (2009).

