## Supplementary figures and images for "Antagonistic pleiotropy plays an important role in governing the evolution and genetic diversity of SARS-CoV-2"

### Supplement Figure 1

#### Fig S1. Mutation spectra of synonymous changes in different months (2020/3-2021/6)

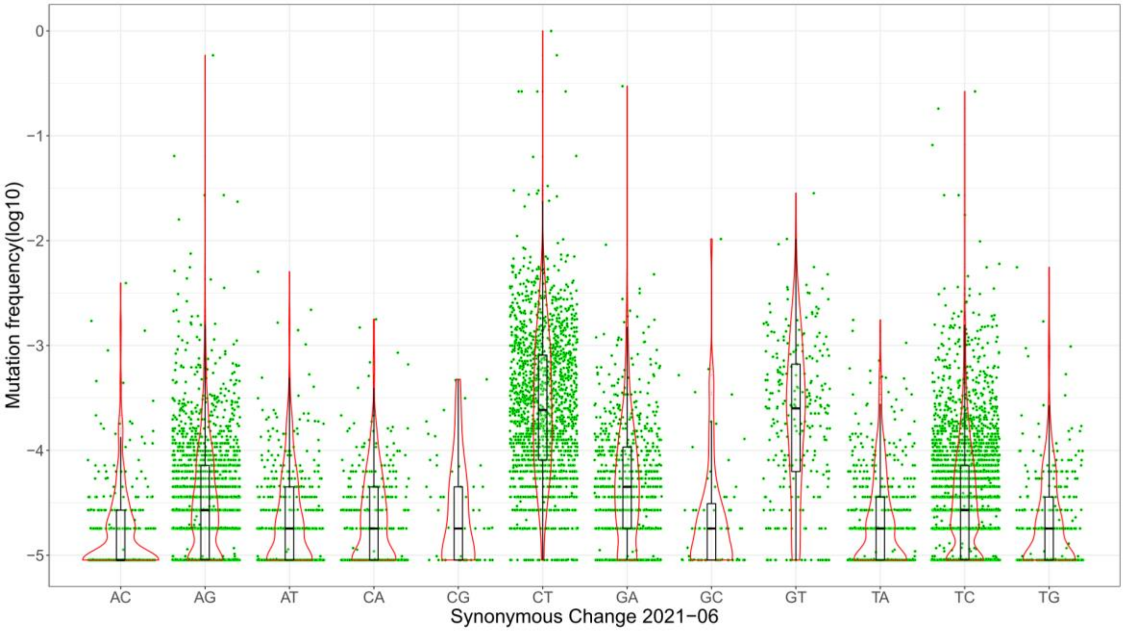


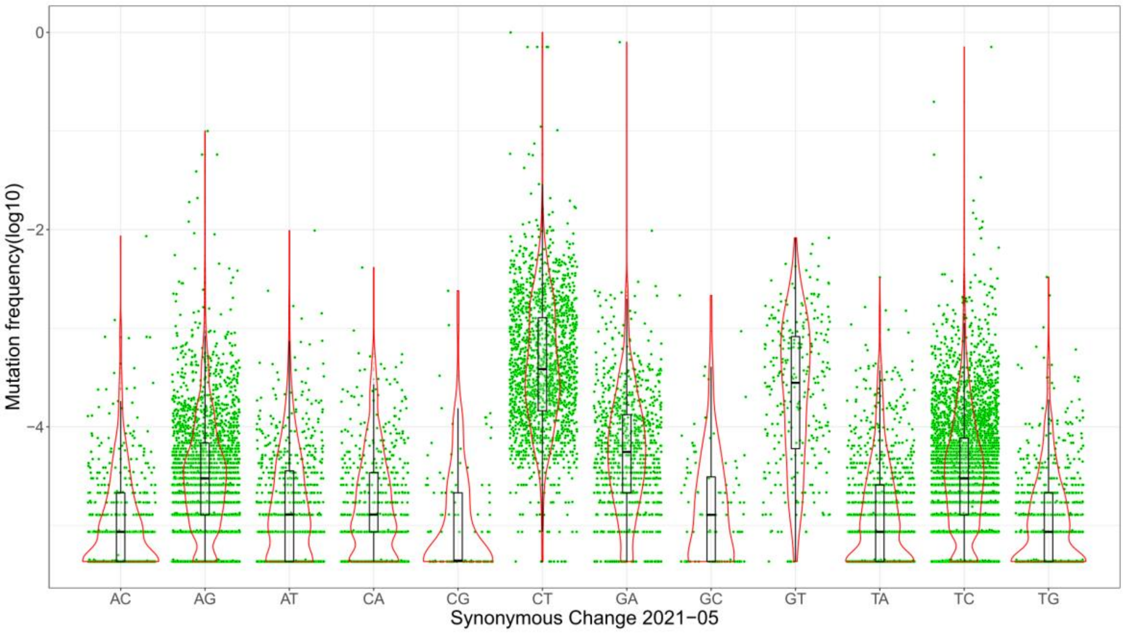

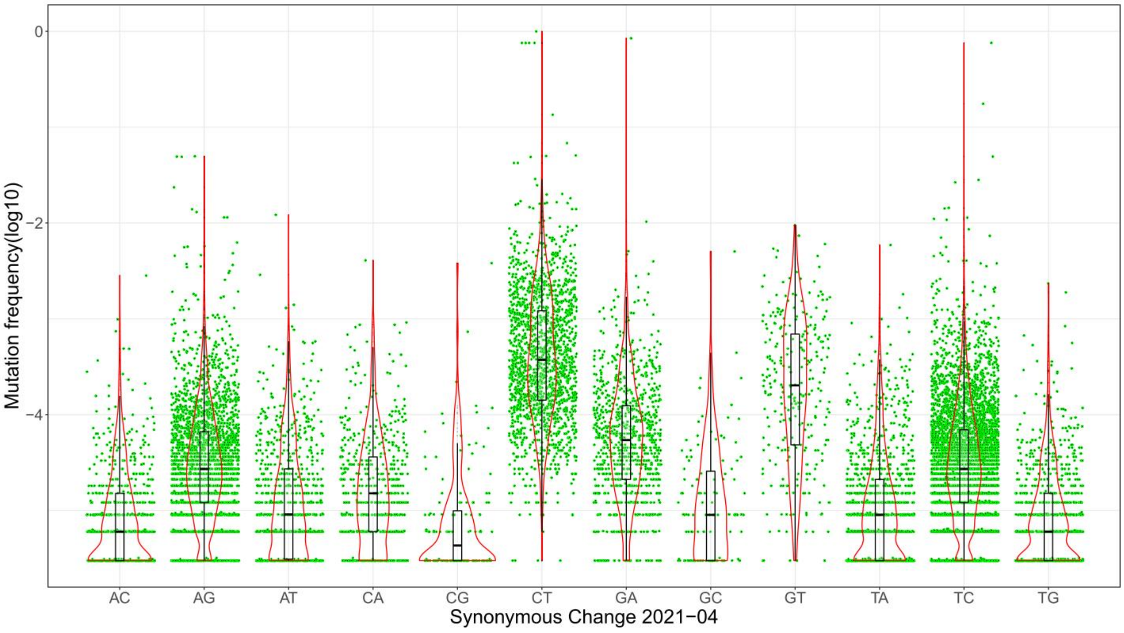

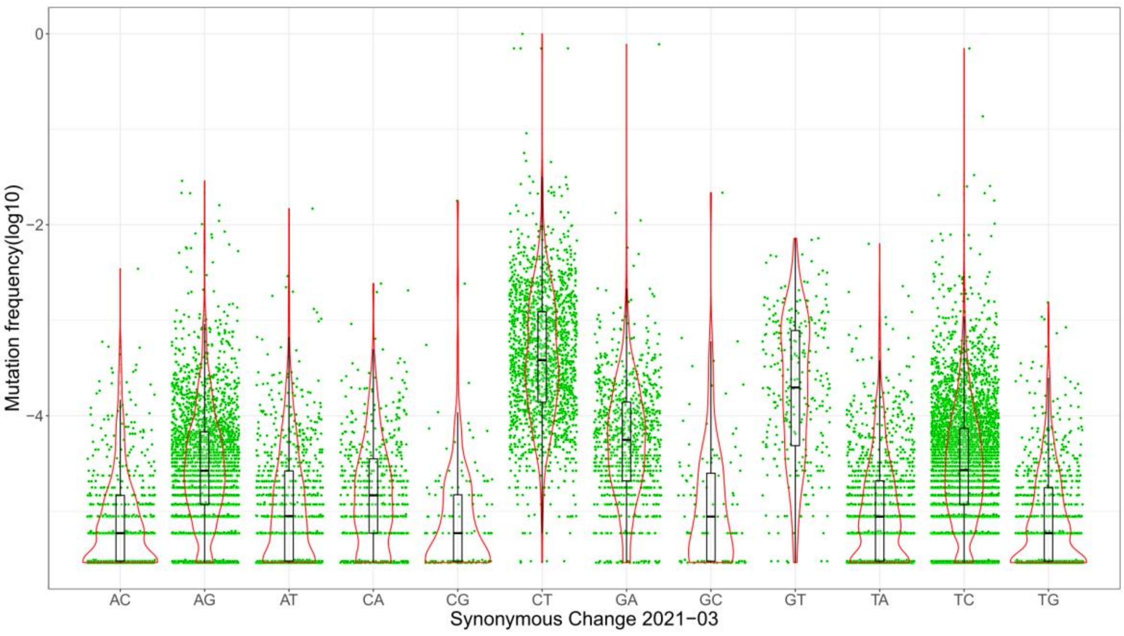

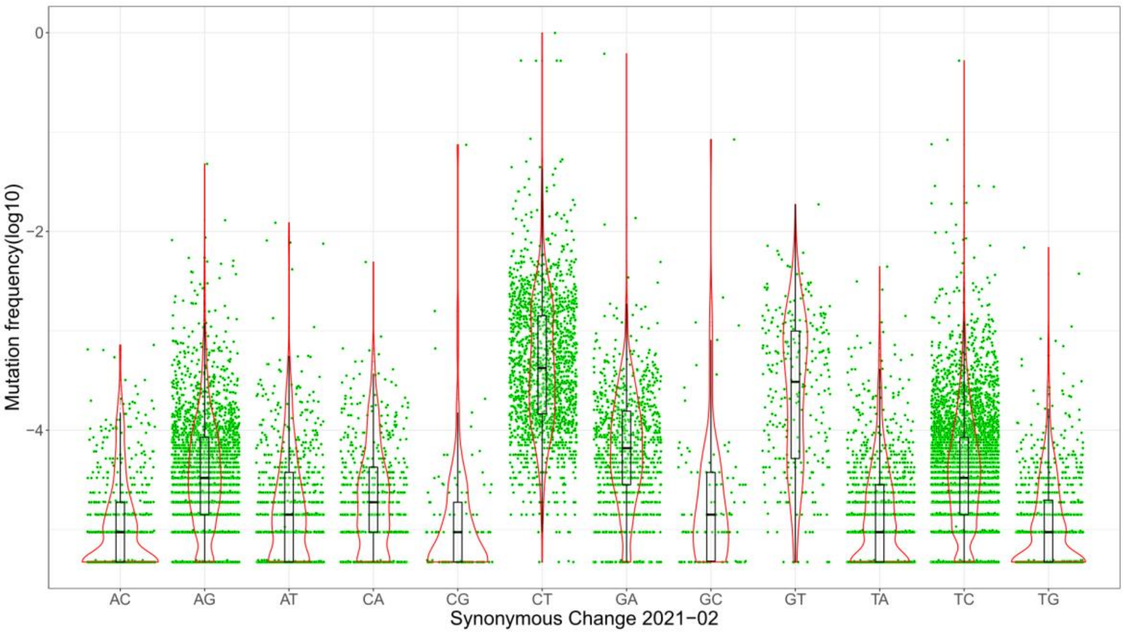

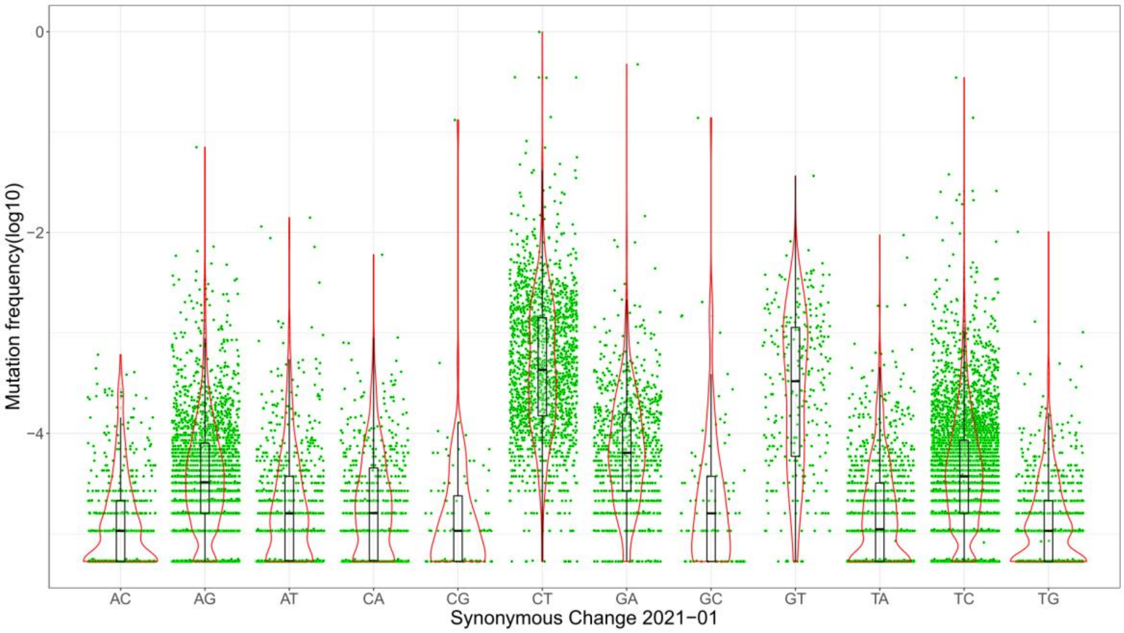

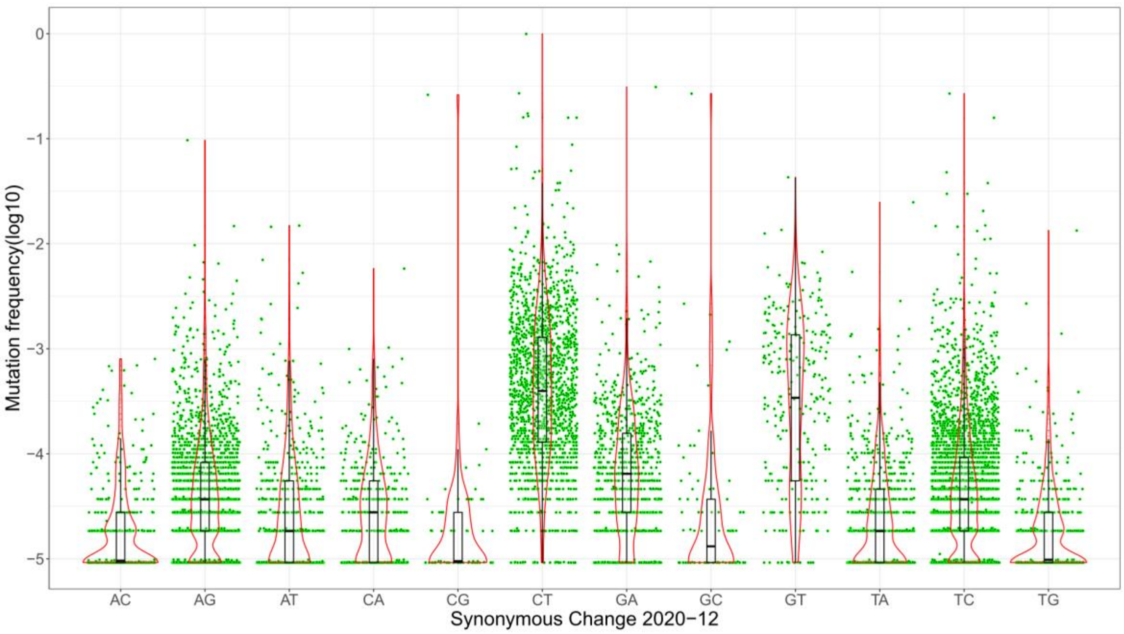

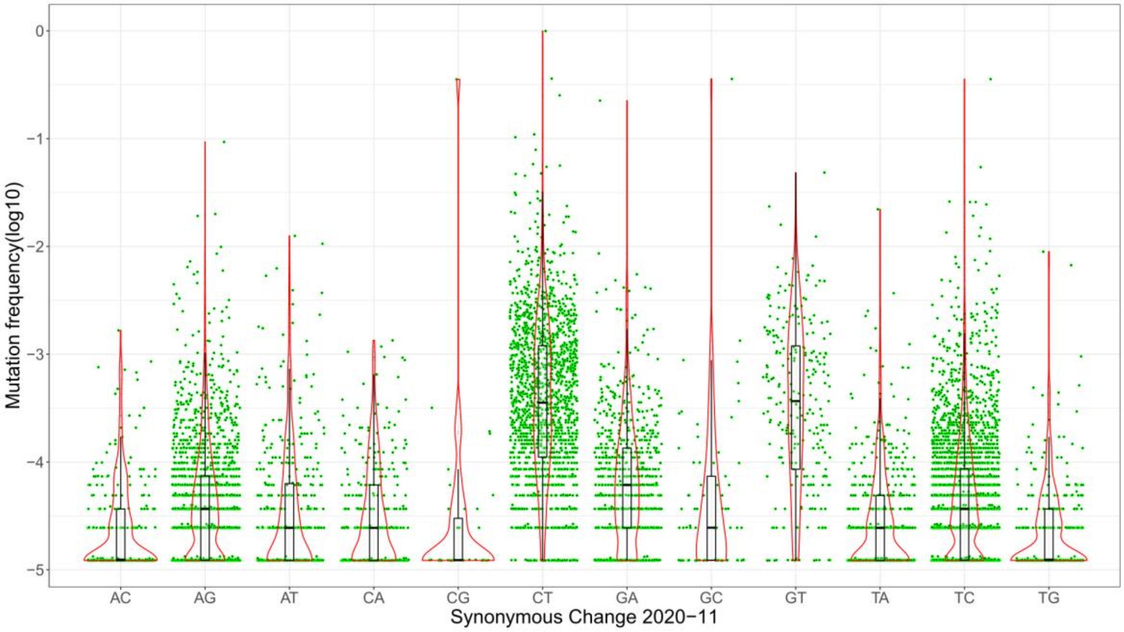

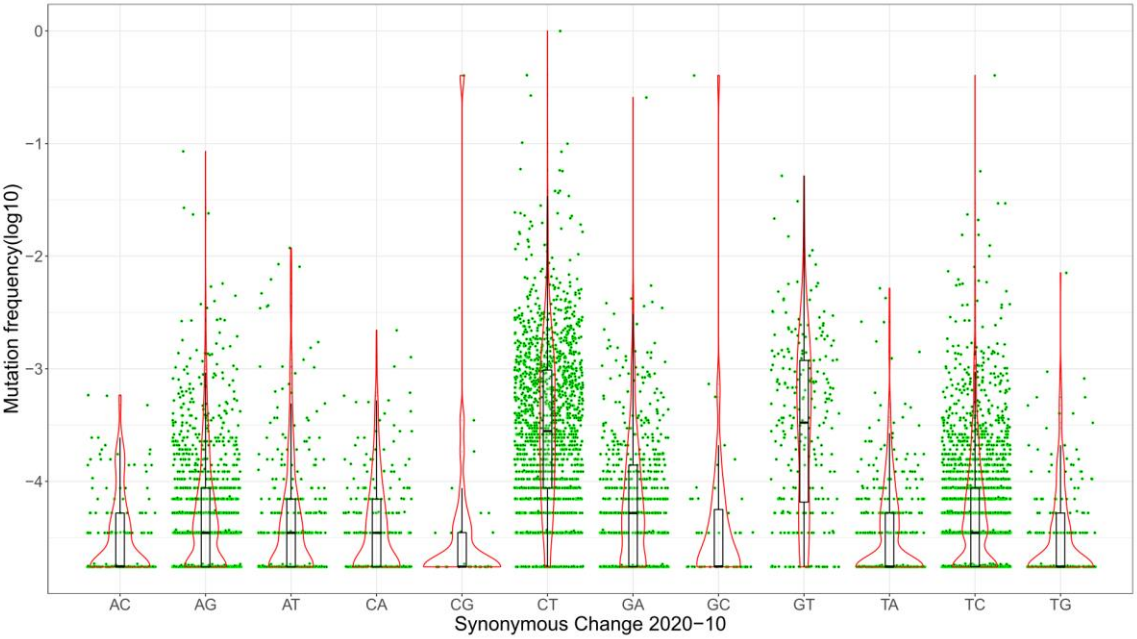

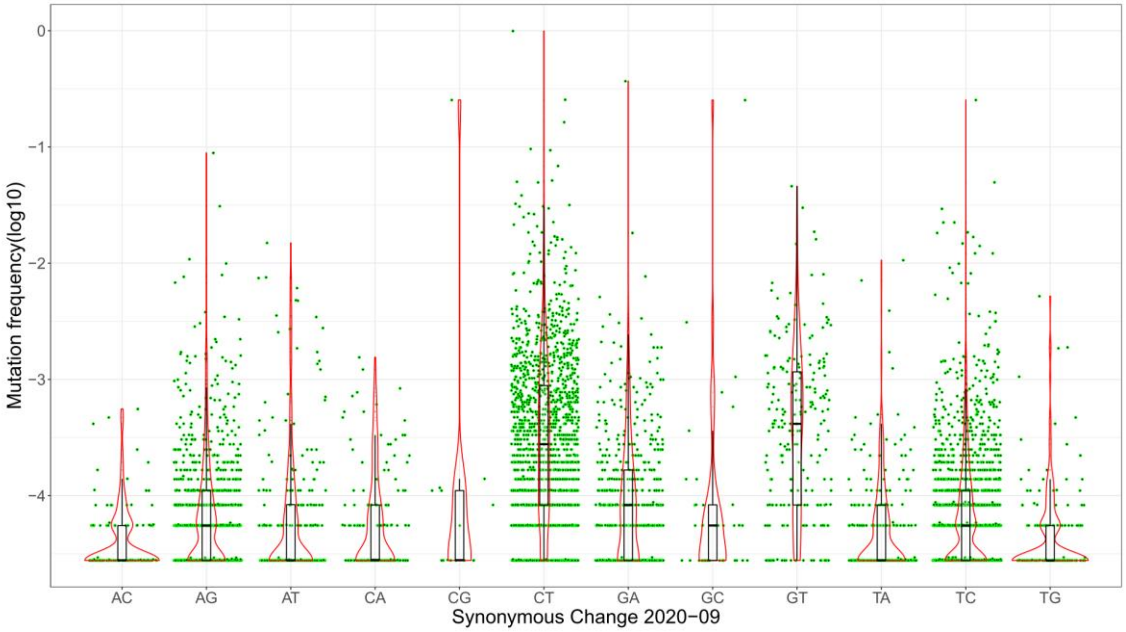

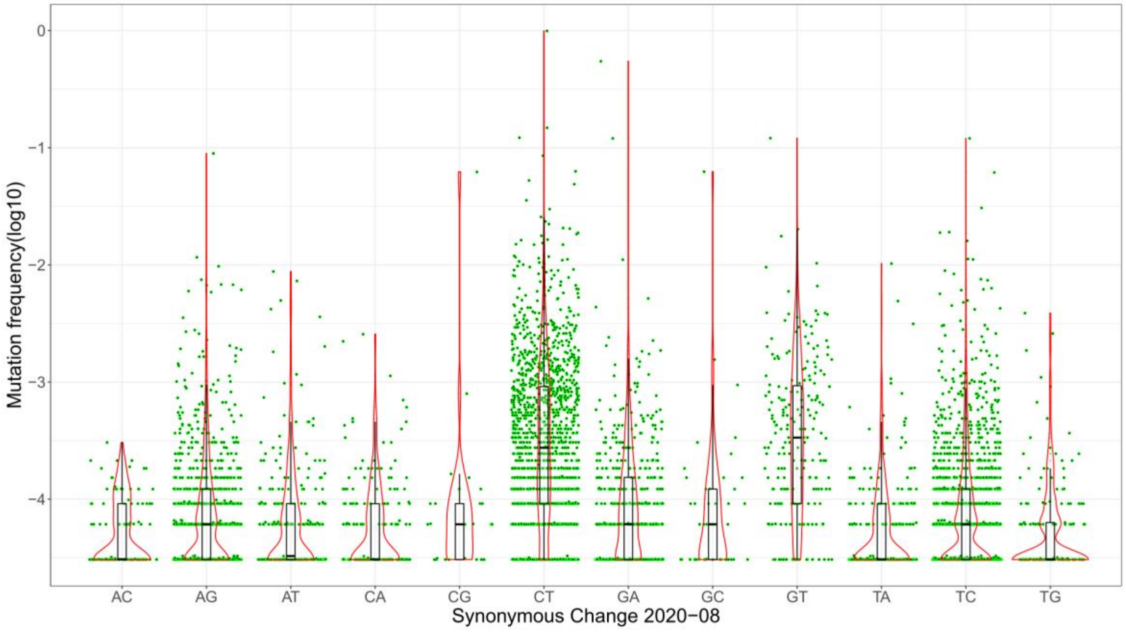

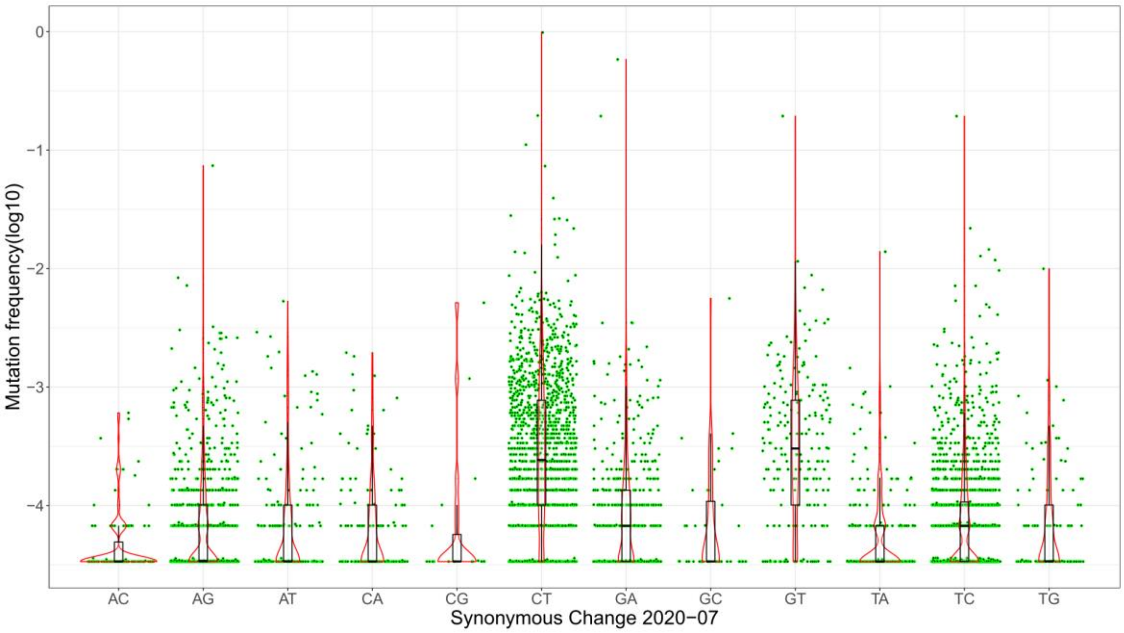

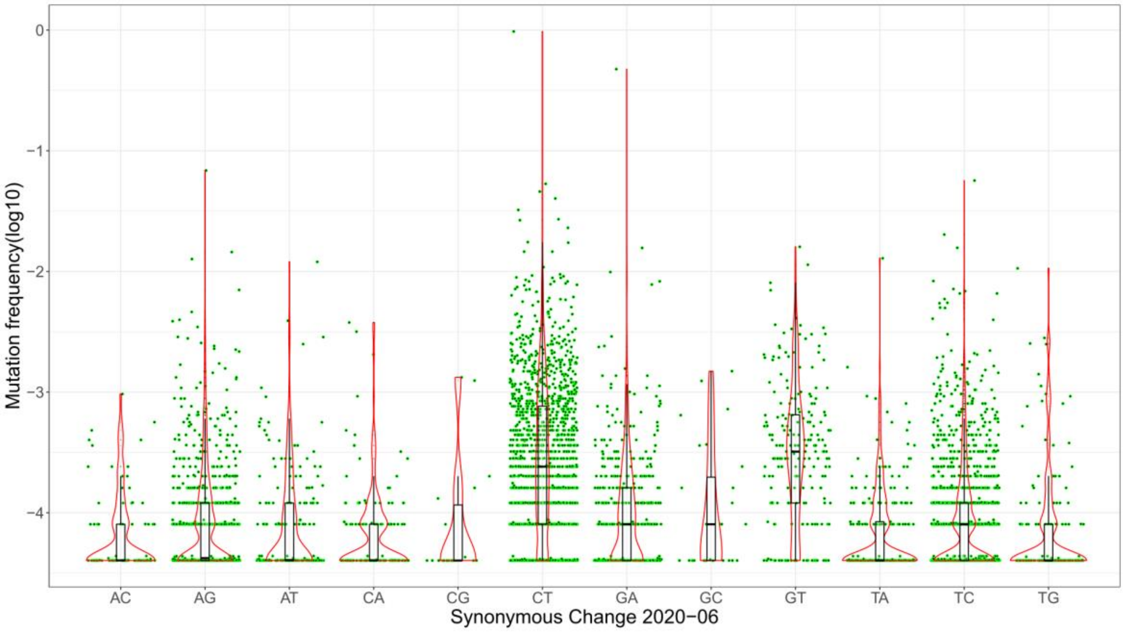

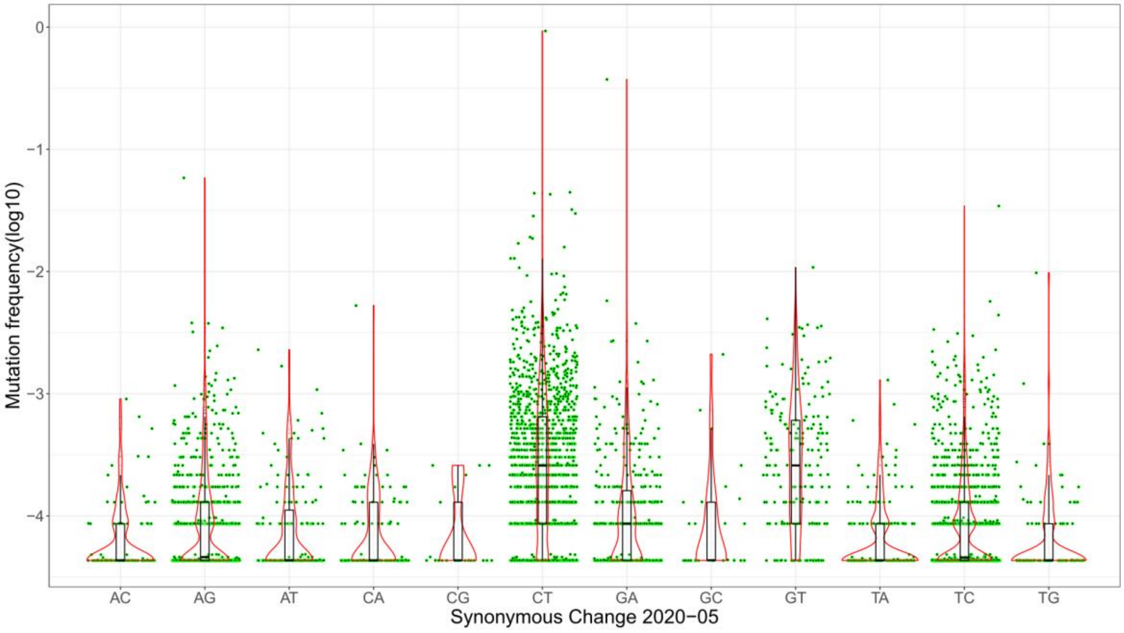

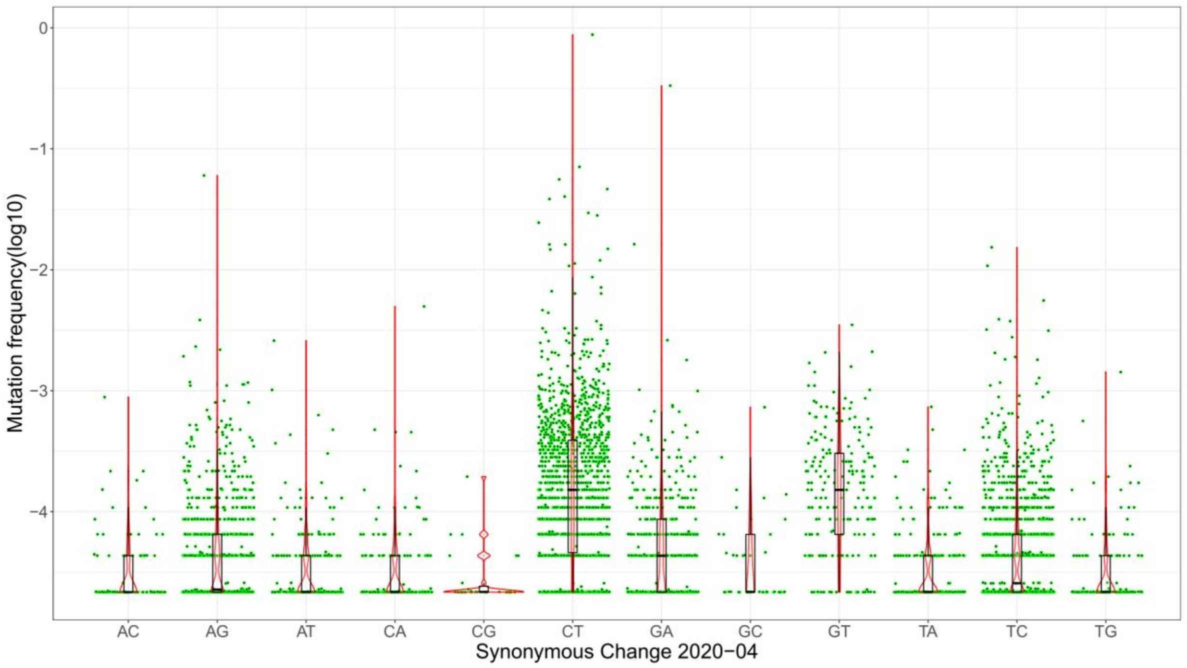

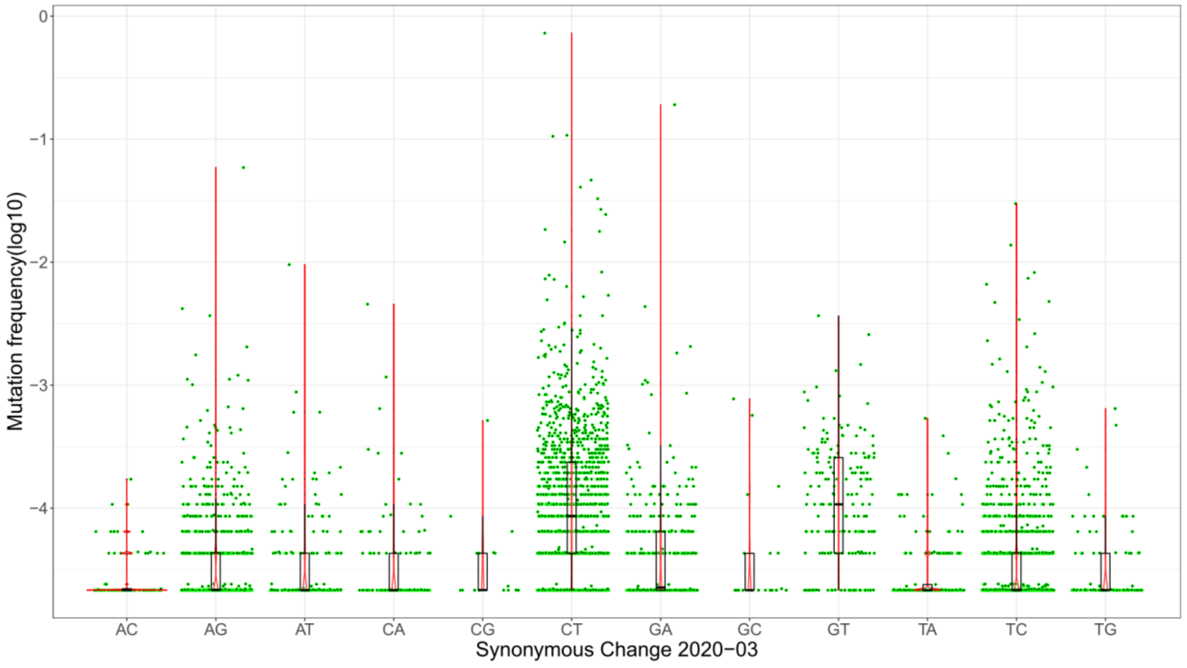

### Supplement Figure 2

**Fig S2. Mutation spectra of nonsynonymous changes in different months (2020/3-2021/6)**

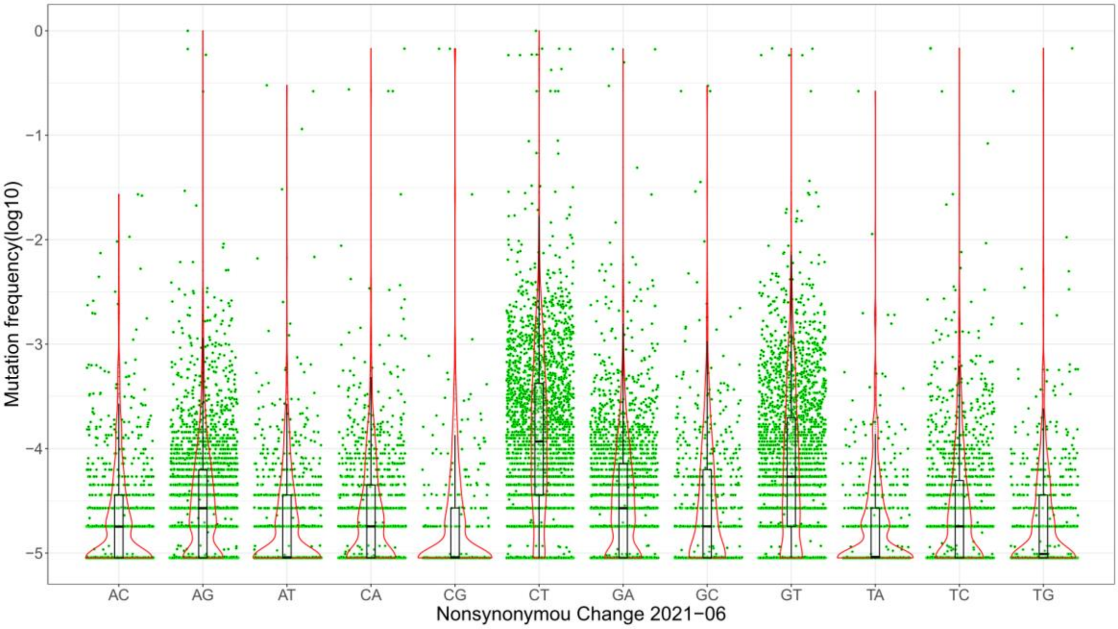


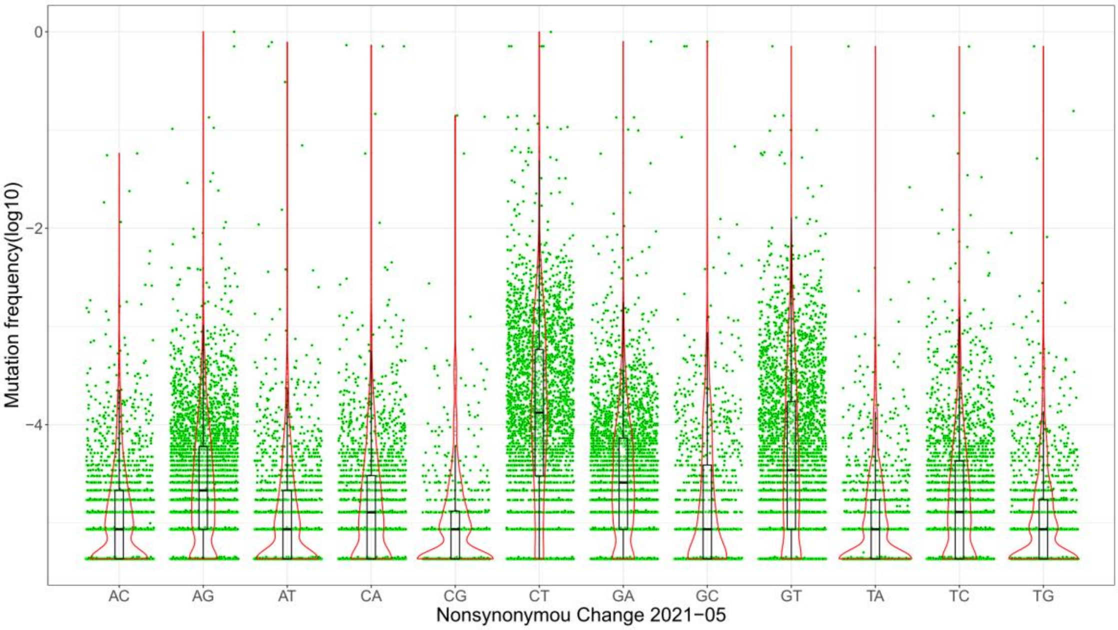

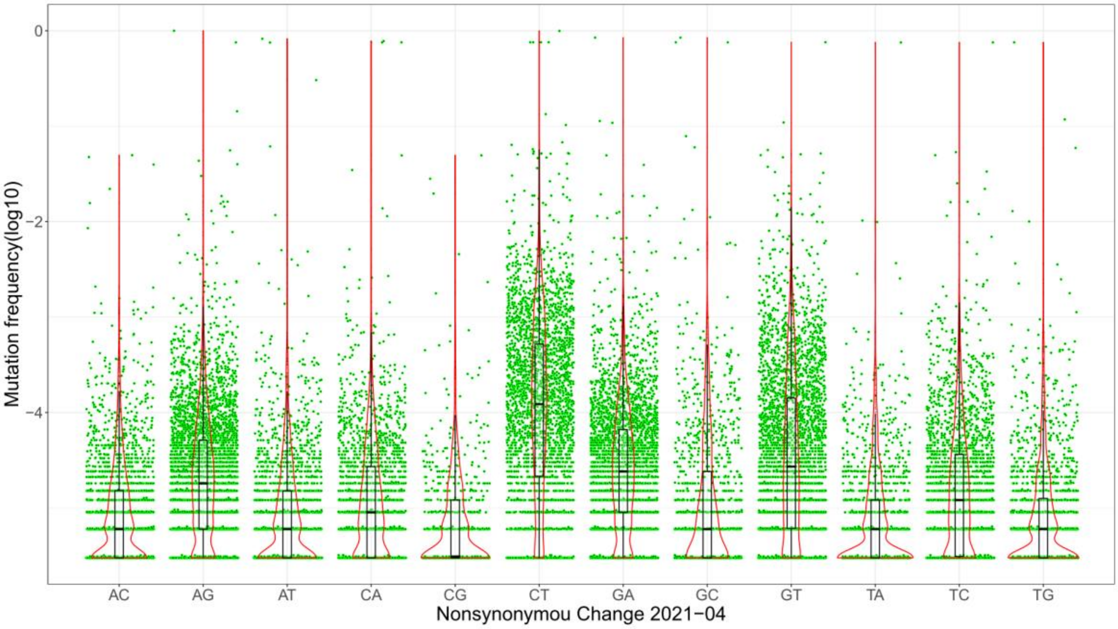

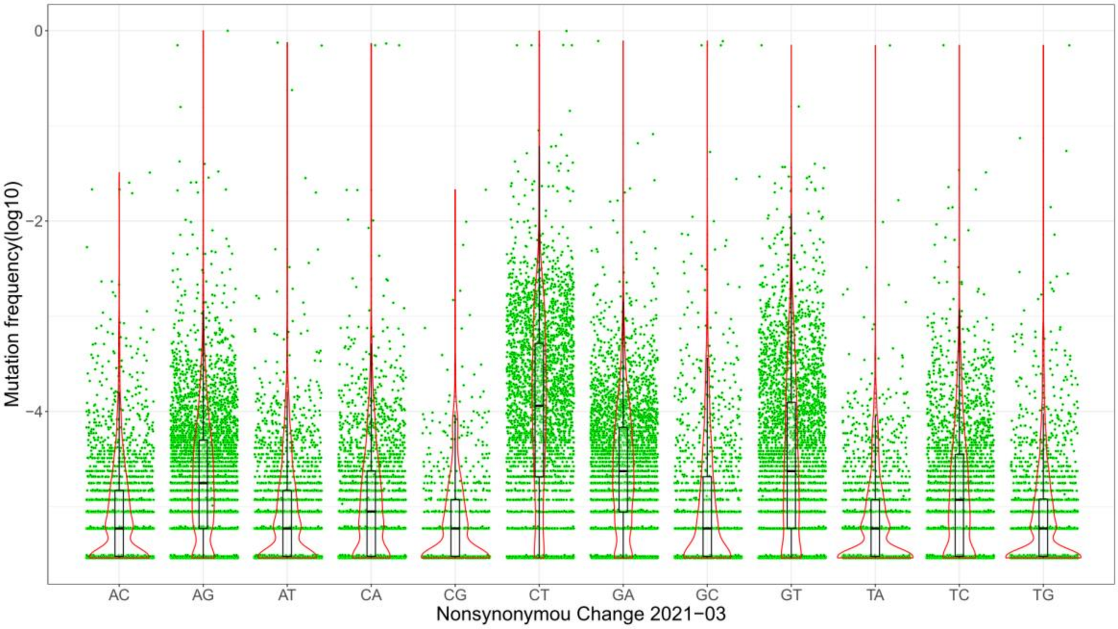

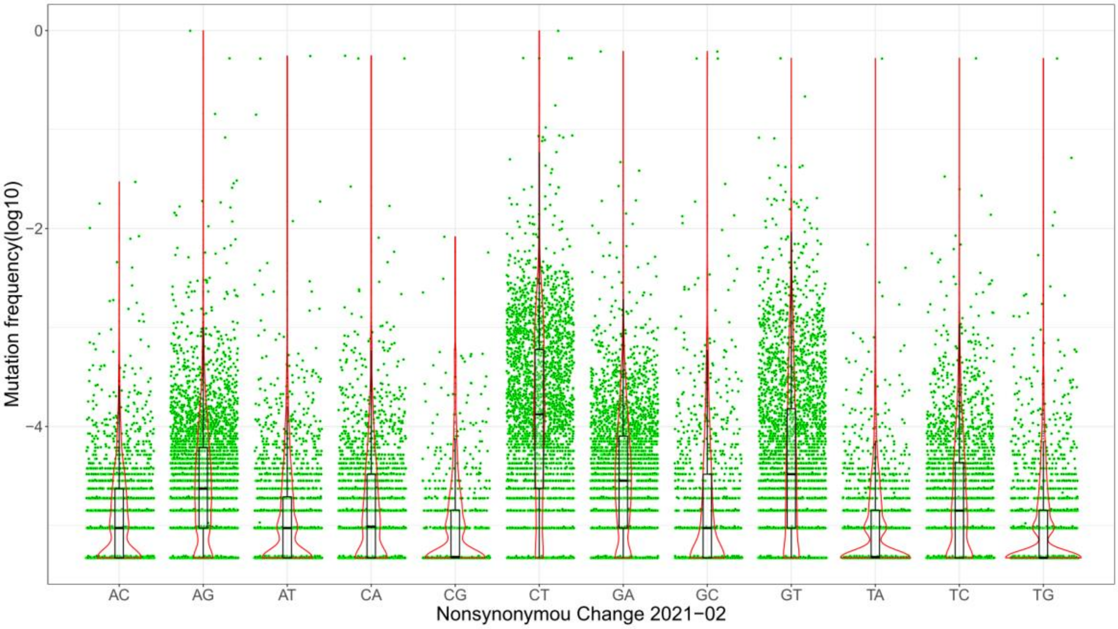

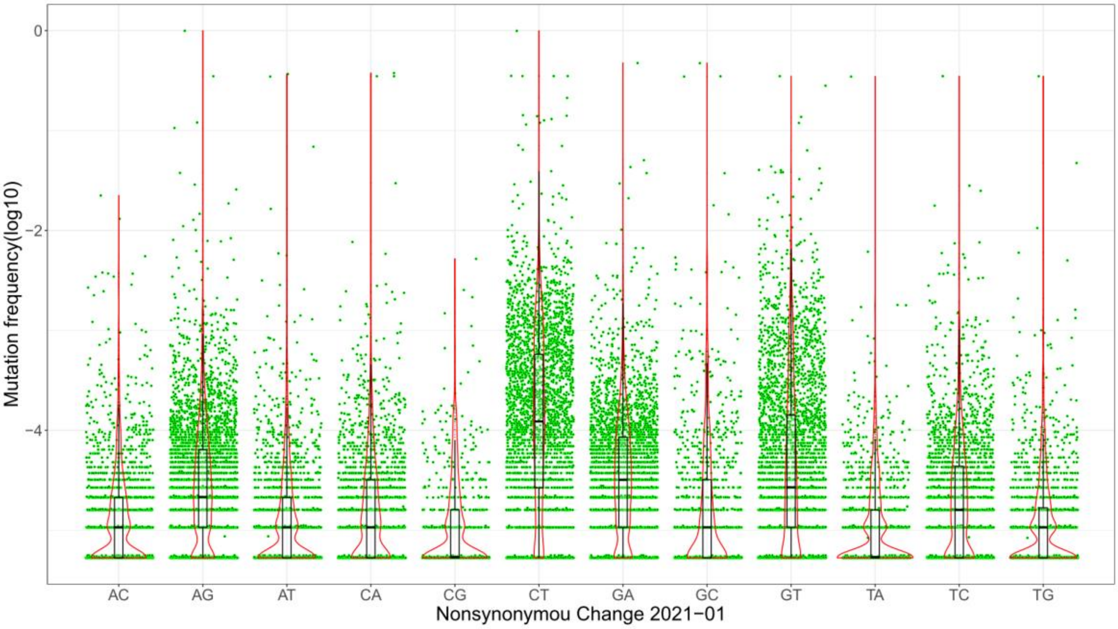

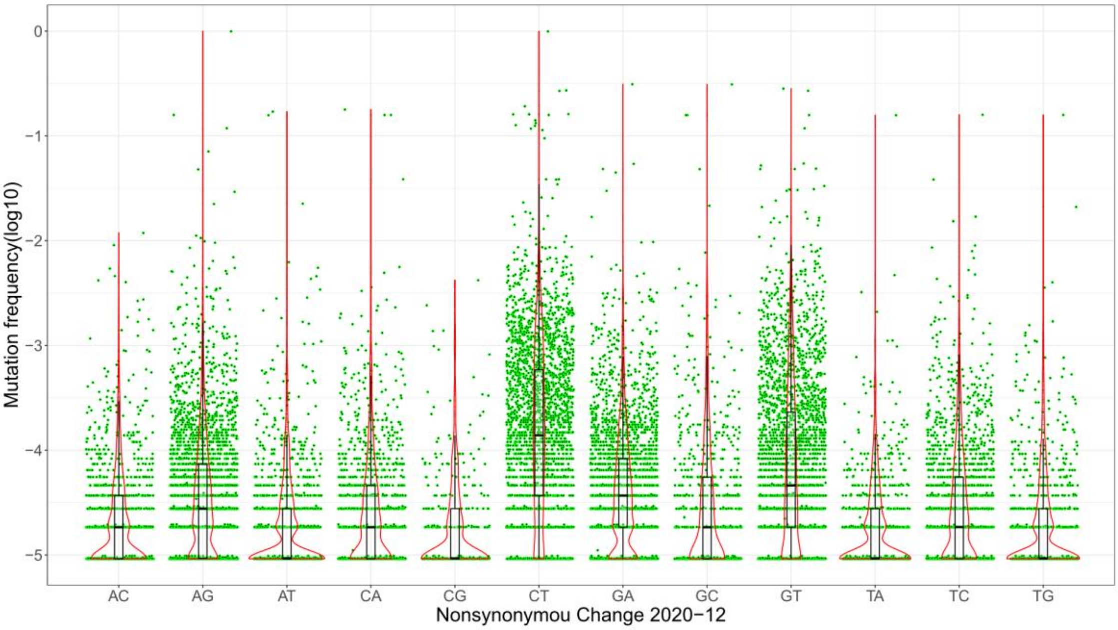

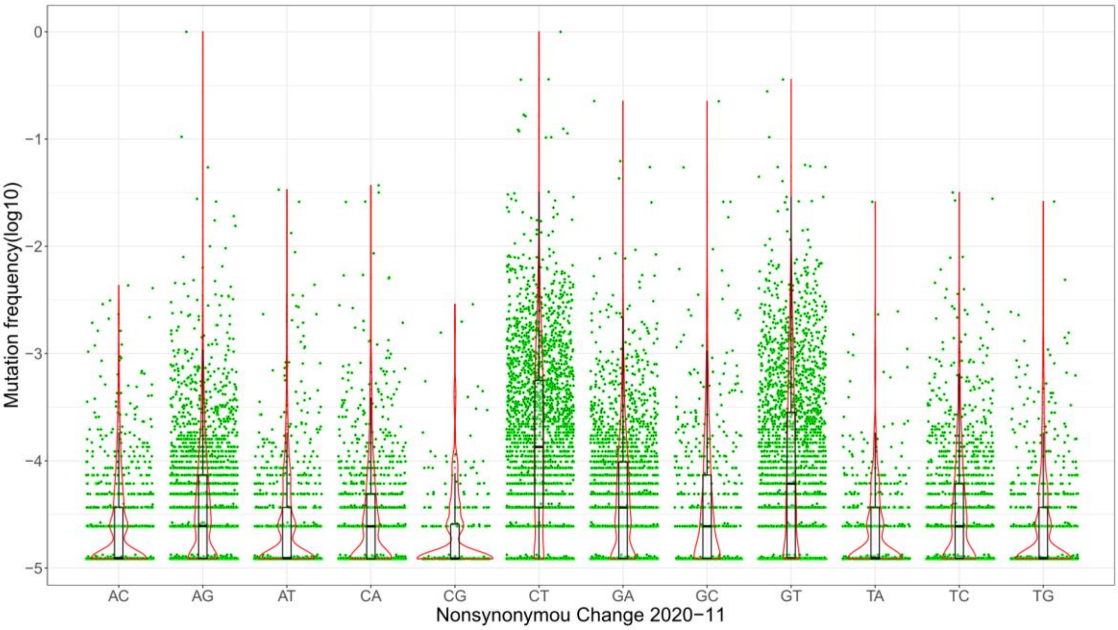

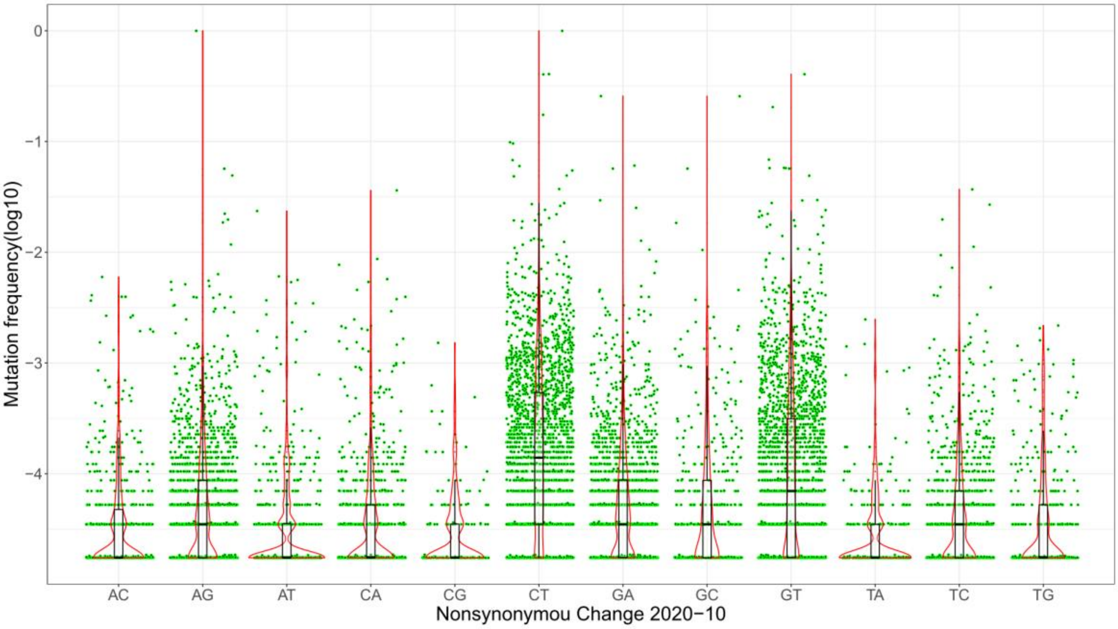

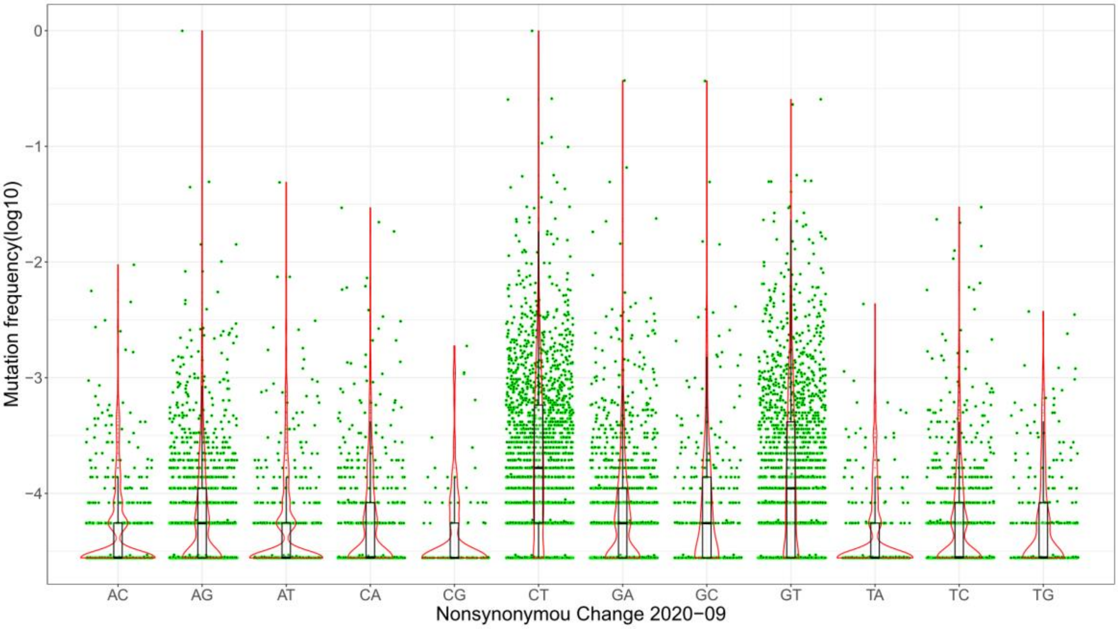

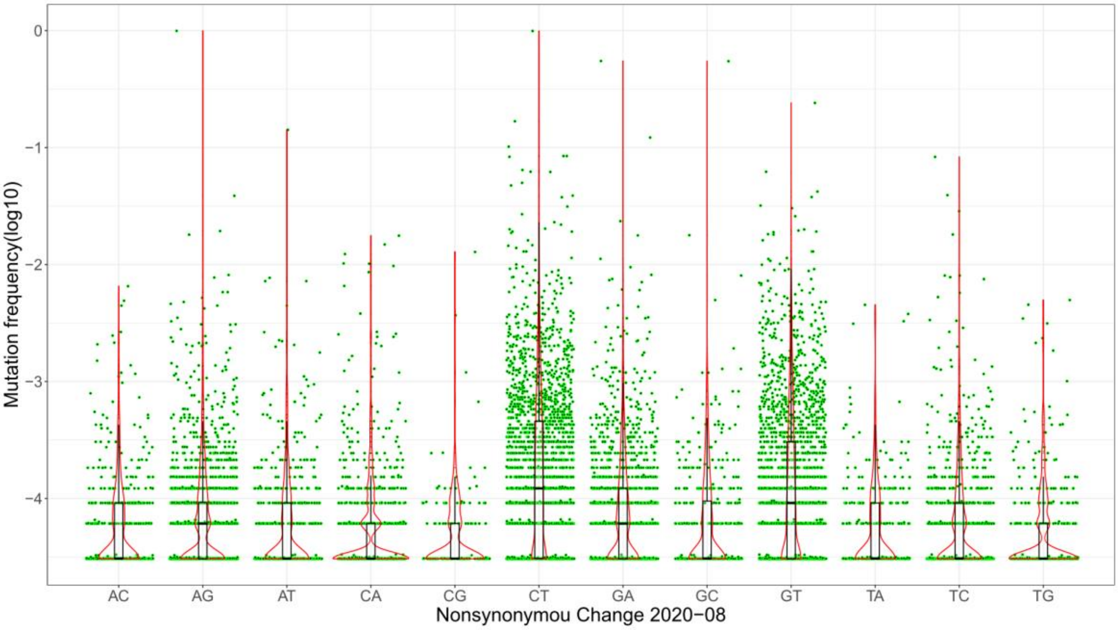

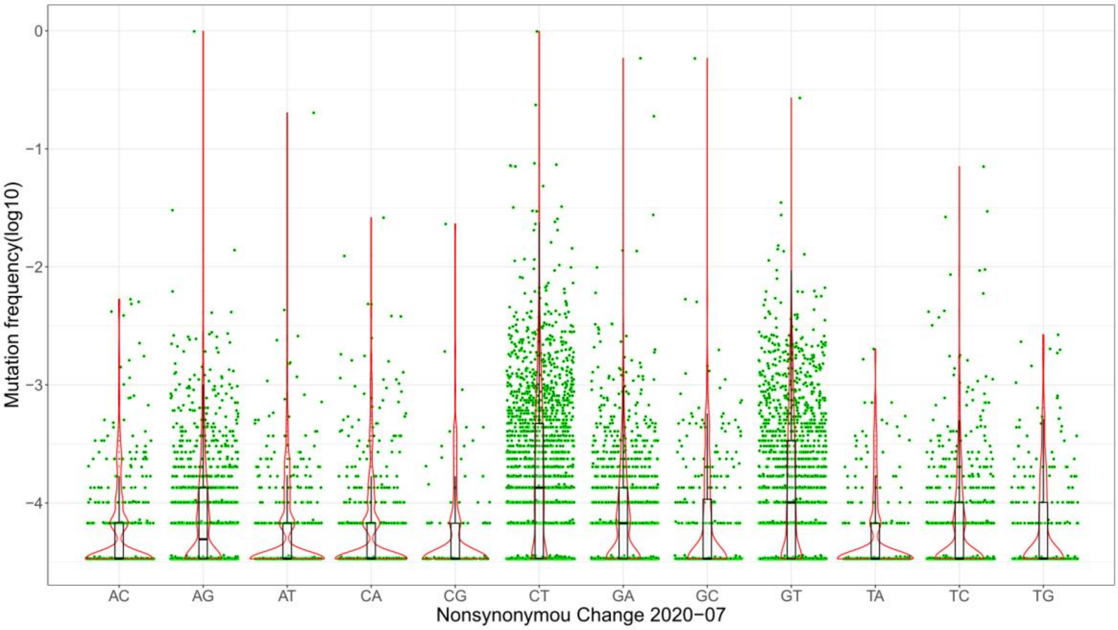

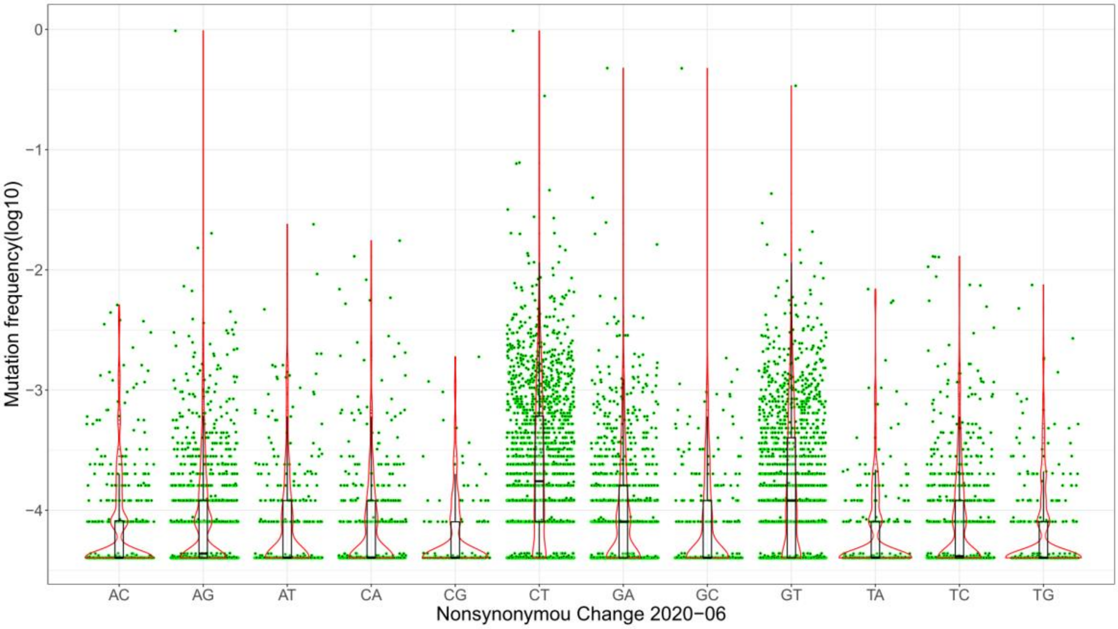

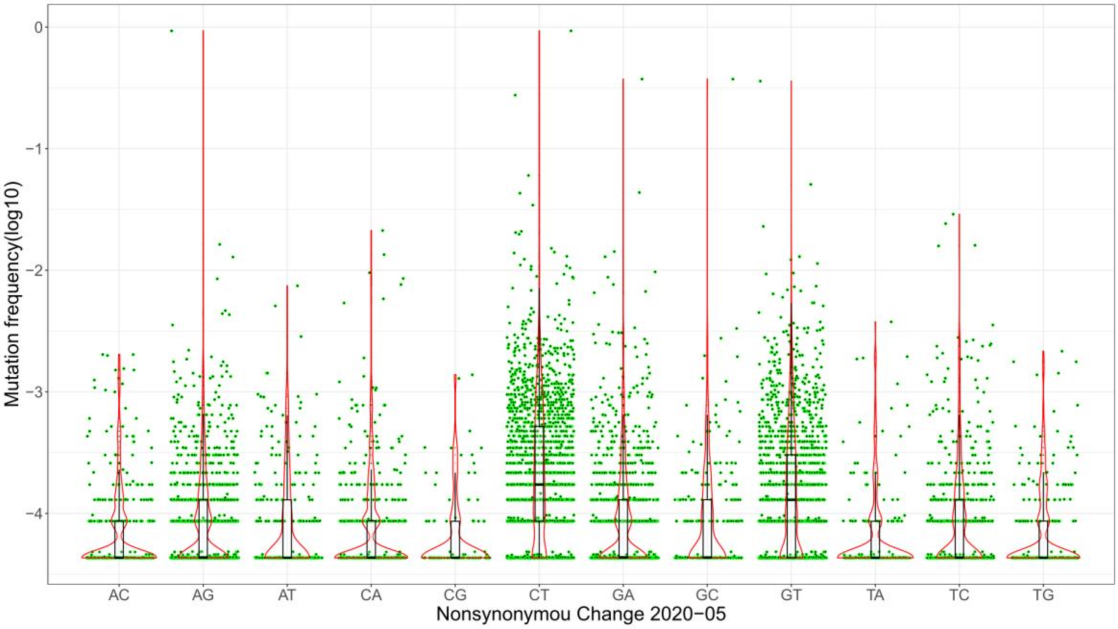

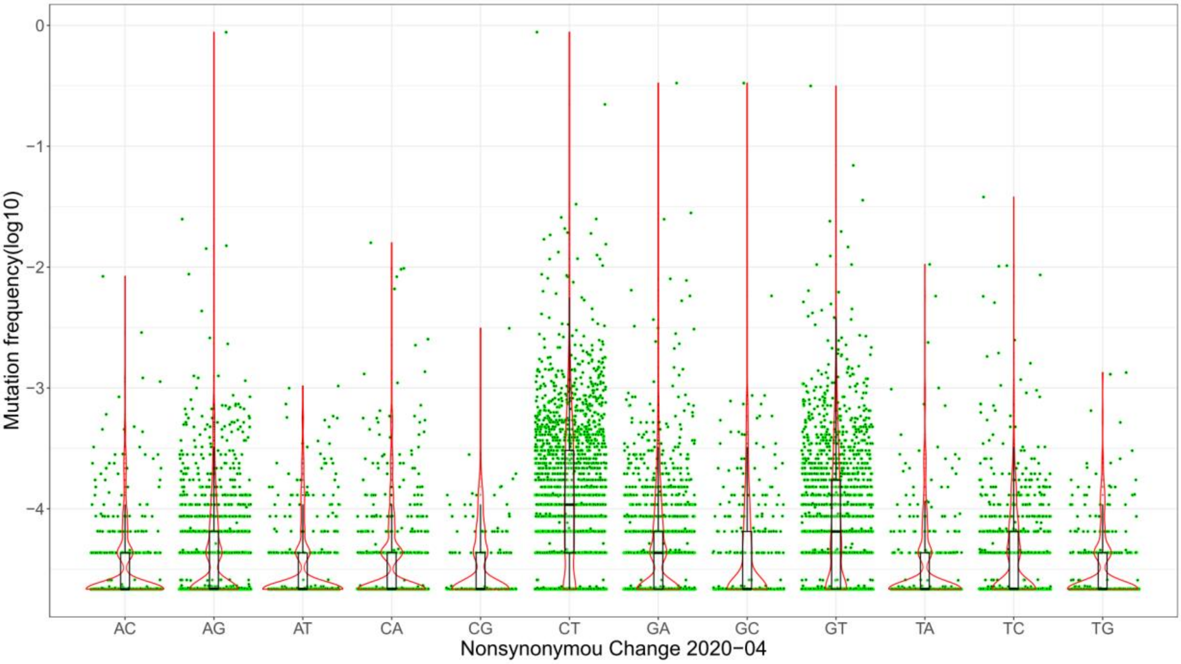

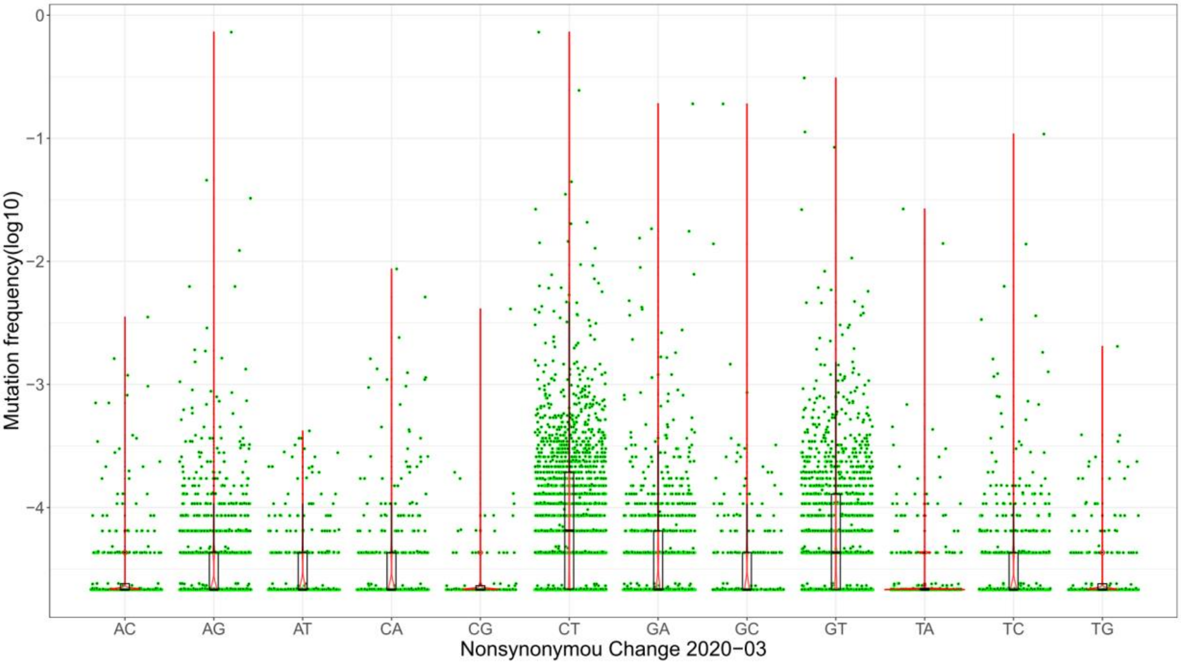

### Supplement Figure 3

**Fig S3. N/S ratios of mutations at different frequencies in different months (2020/3-2021/6)**

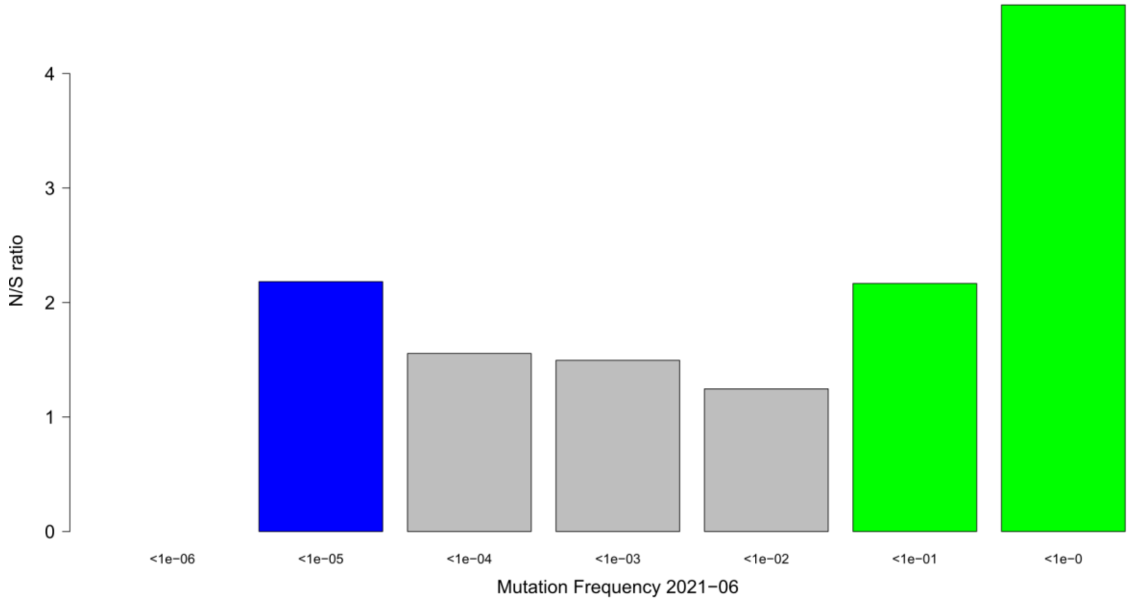


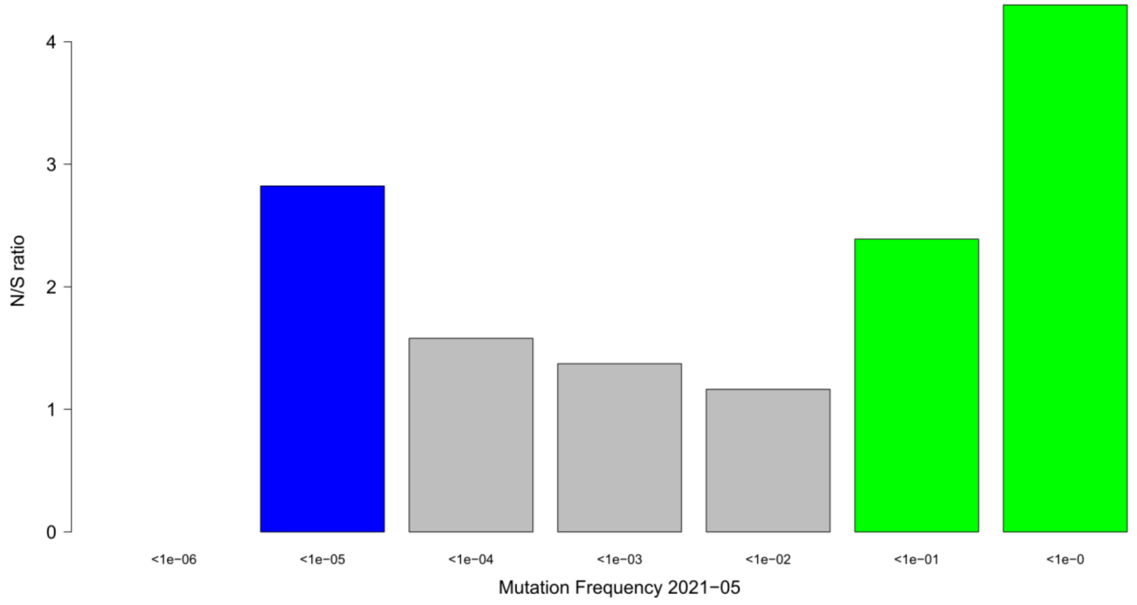

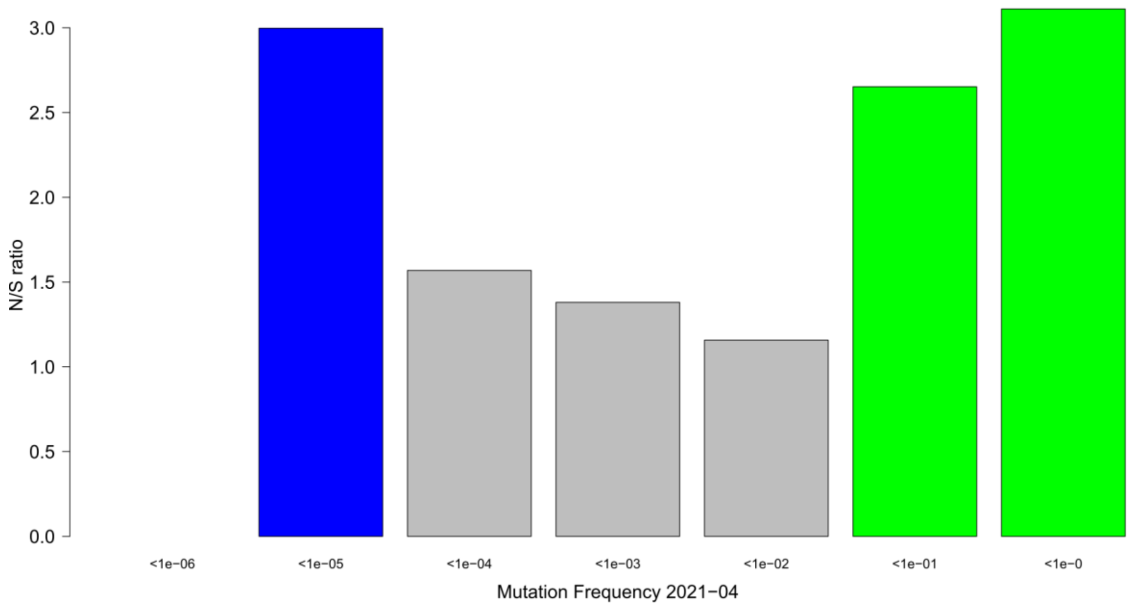

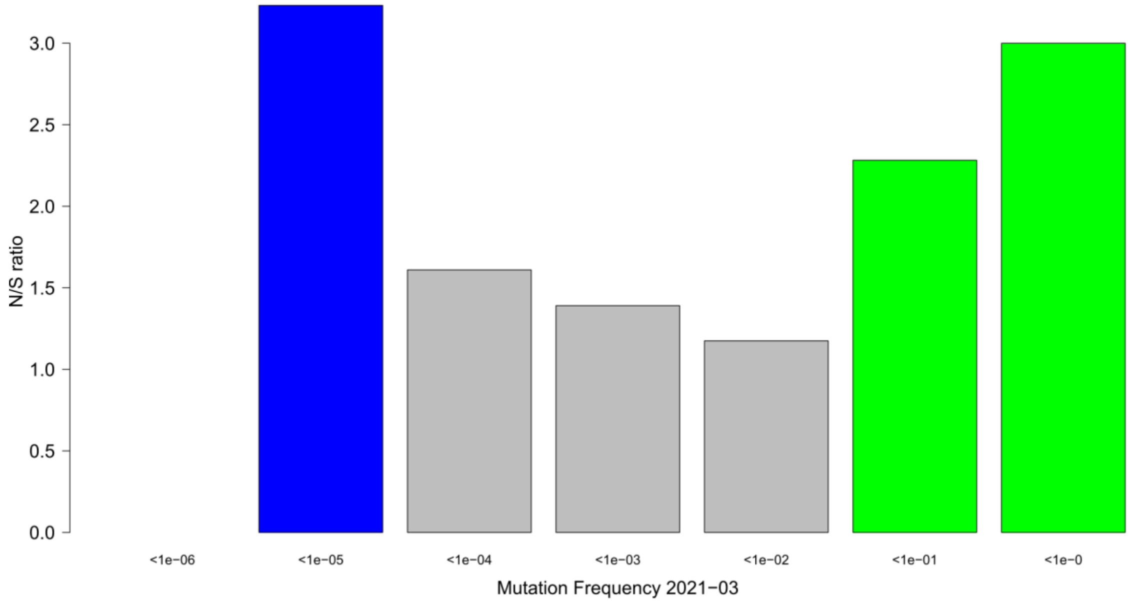

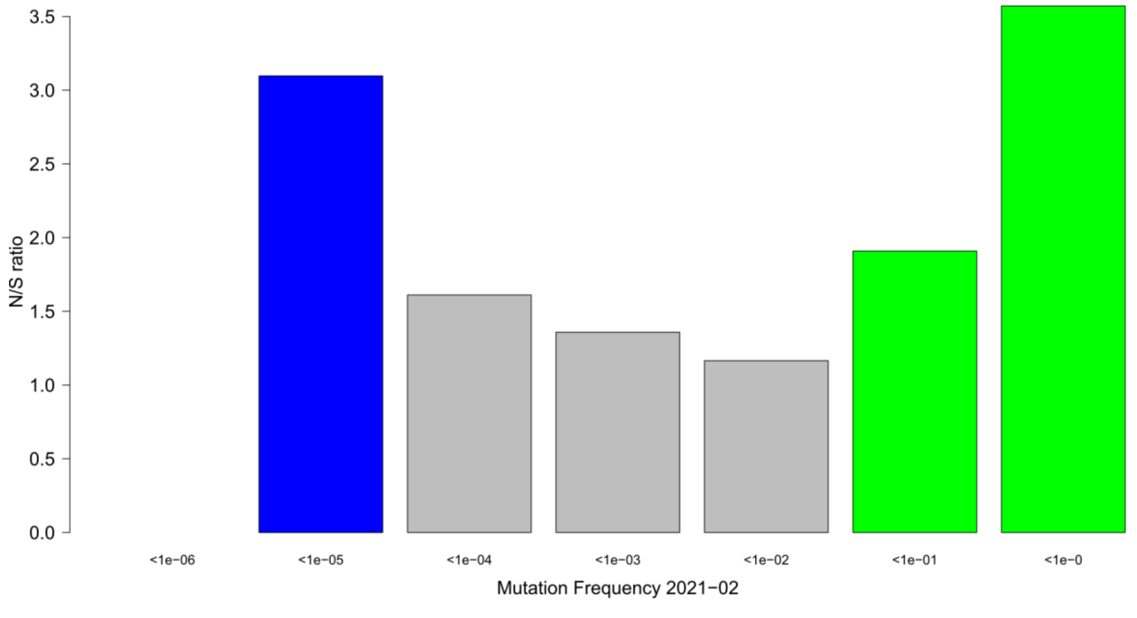

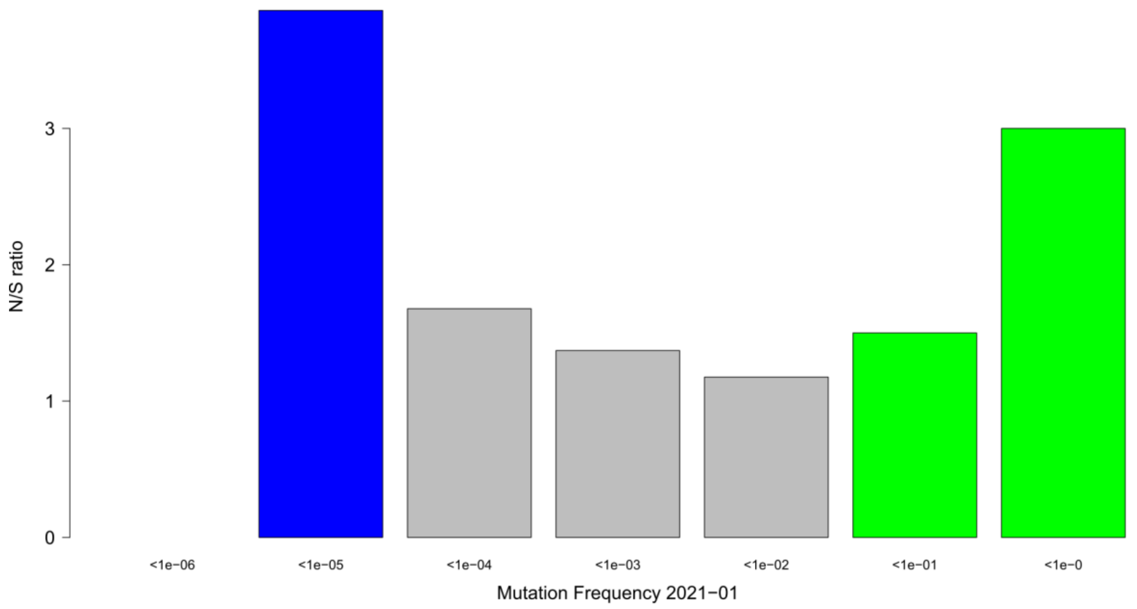

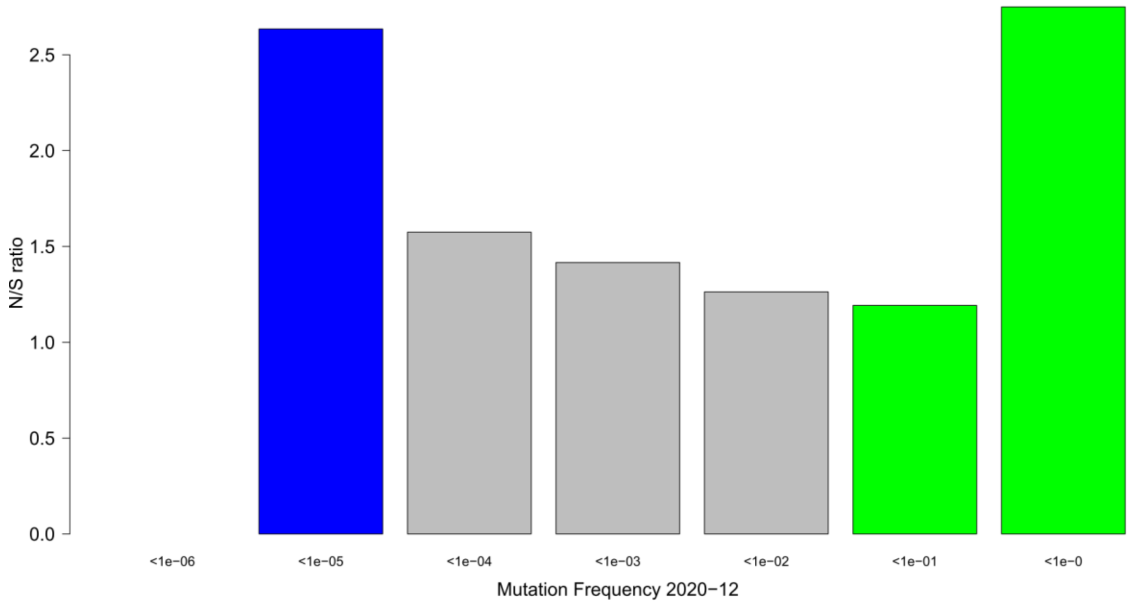

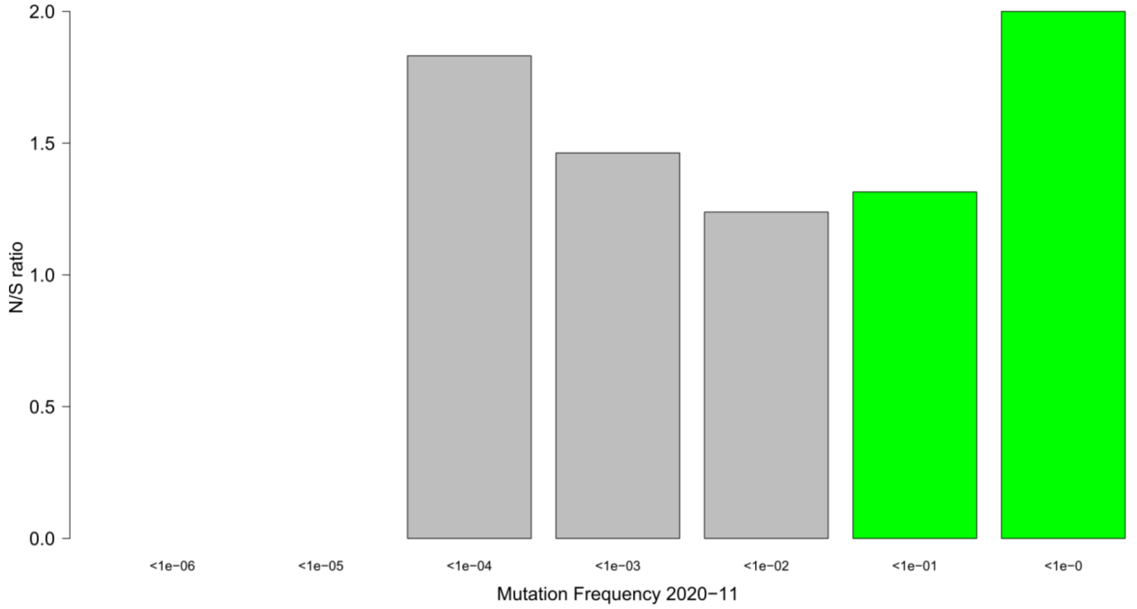

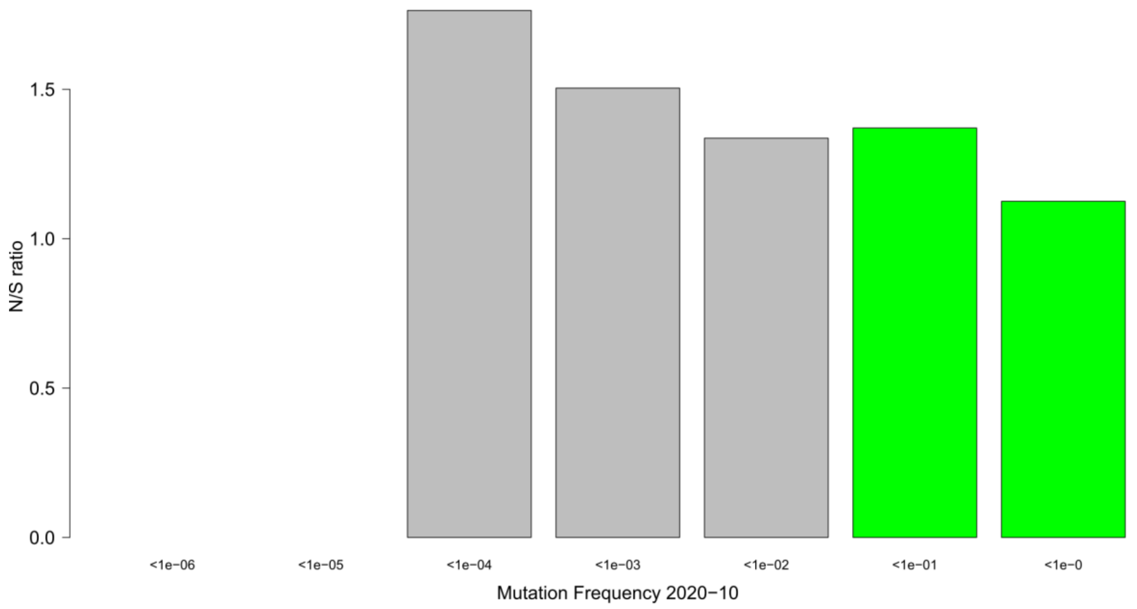

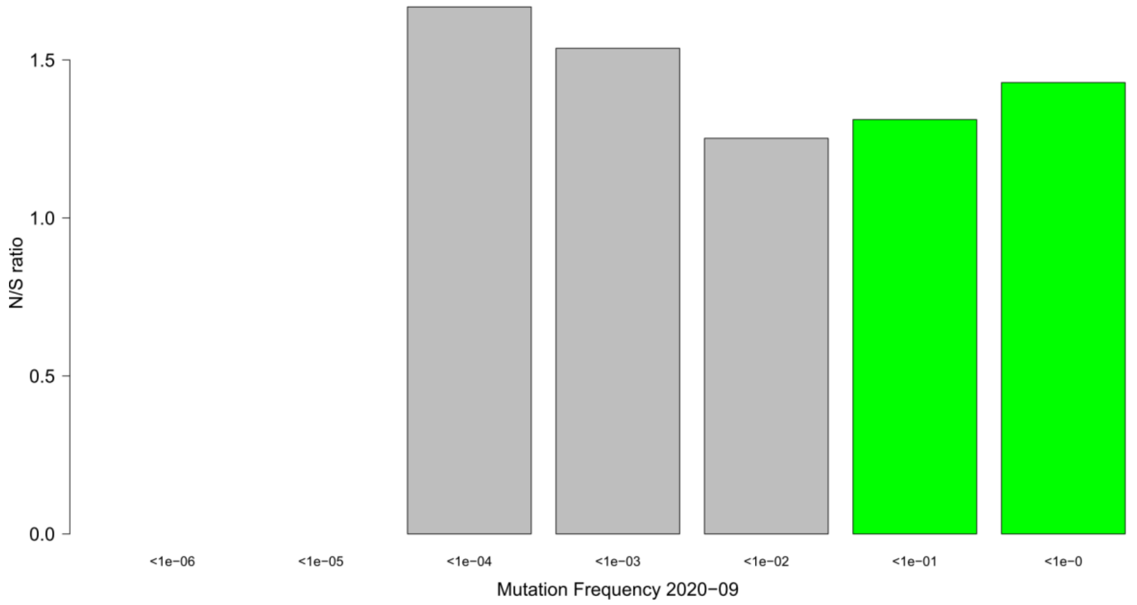

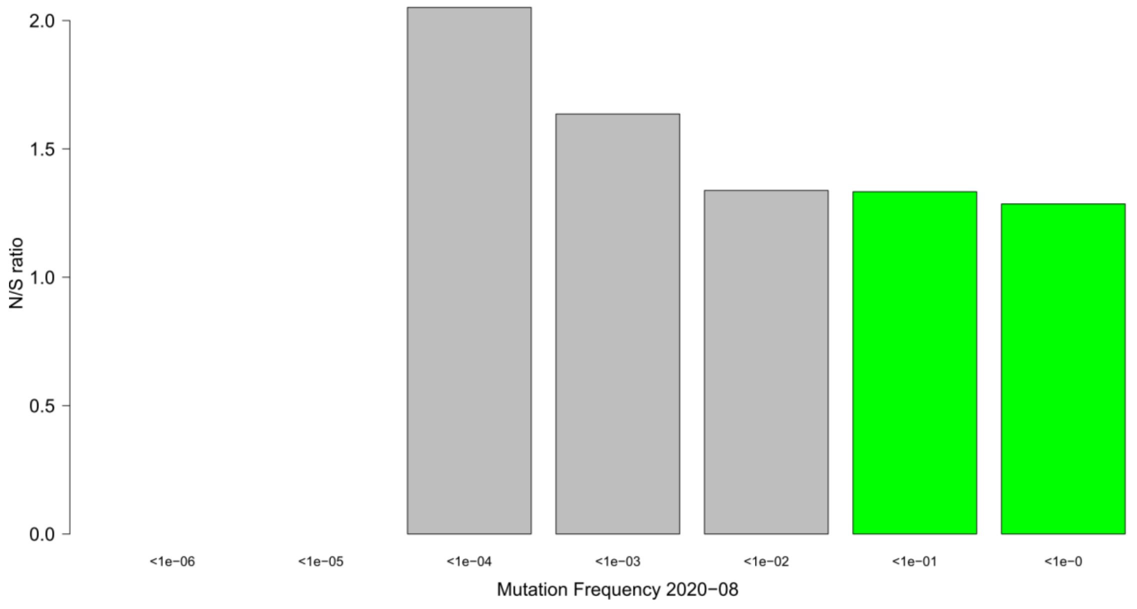

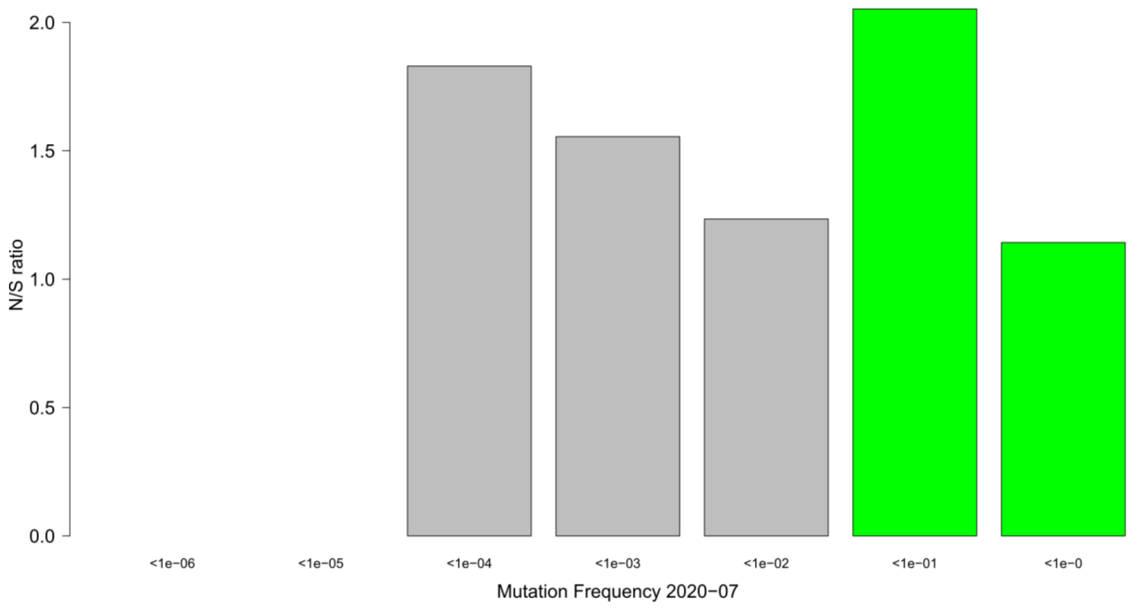

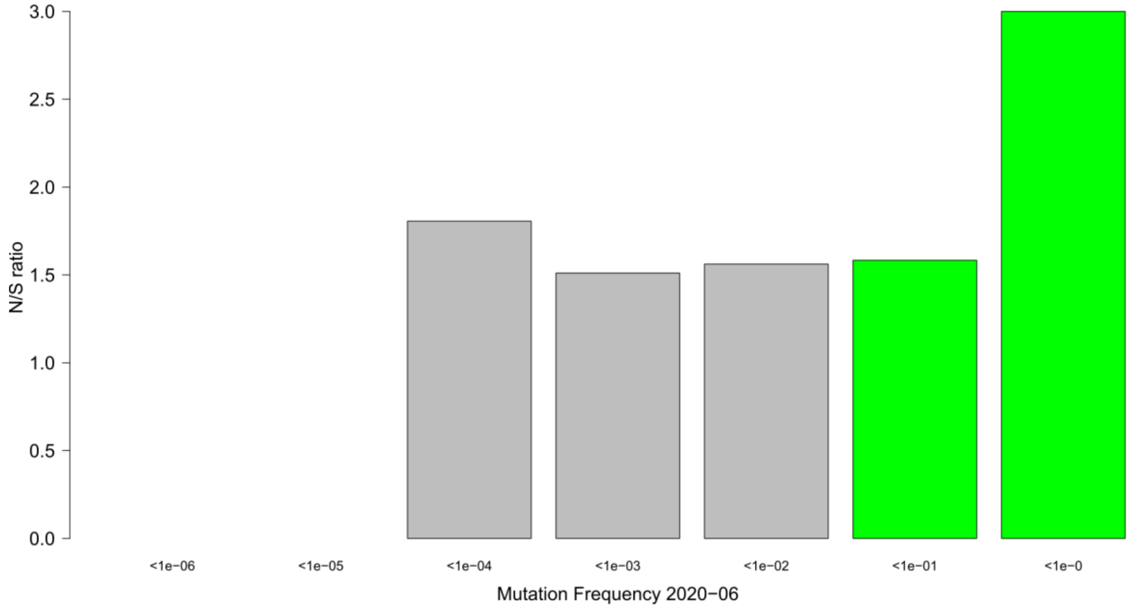

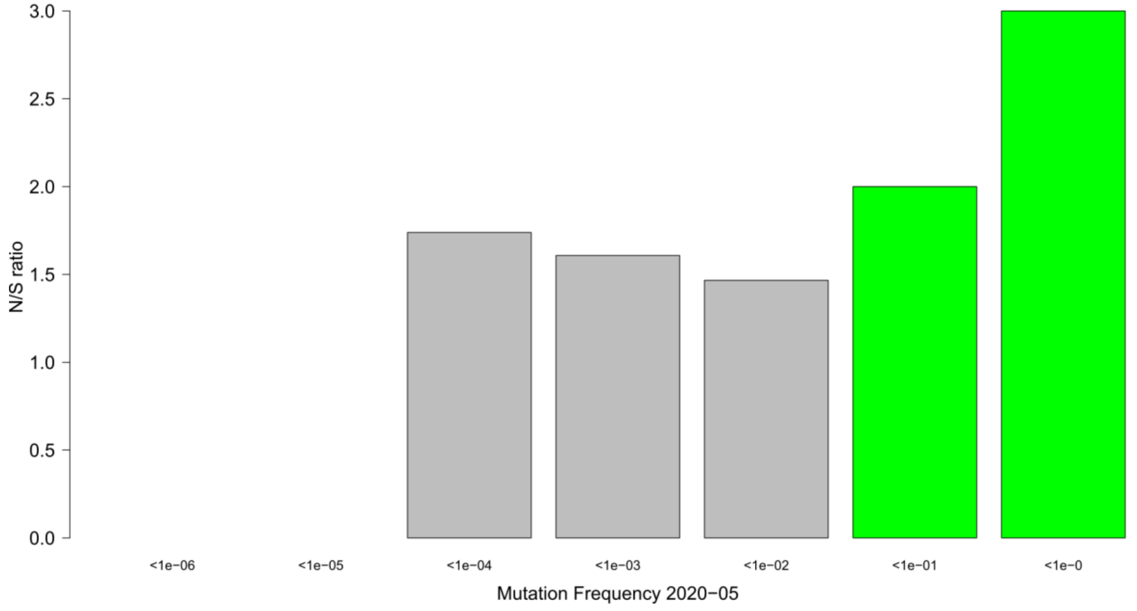

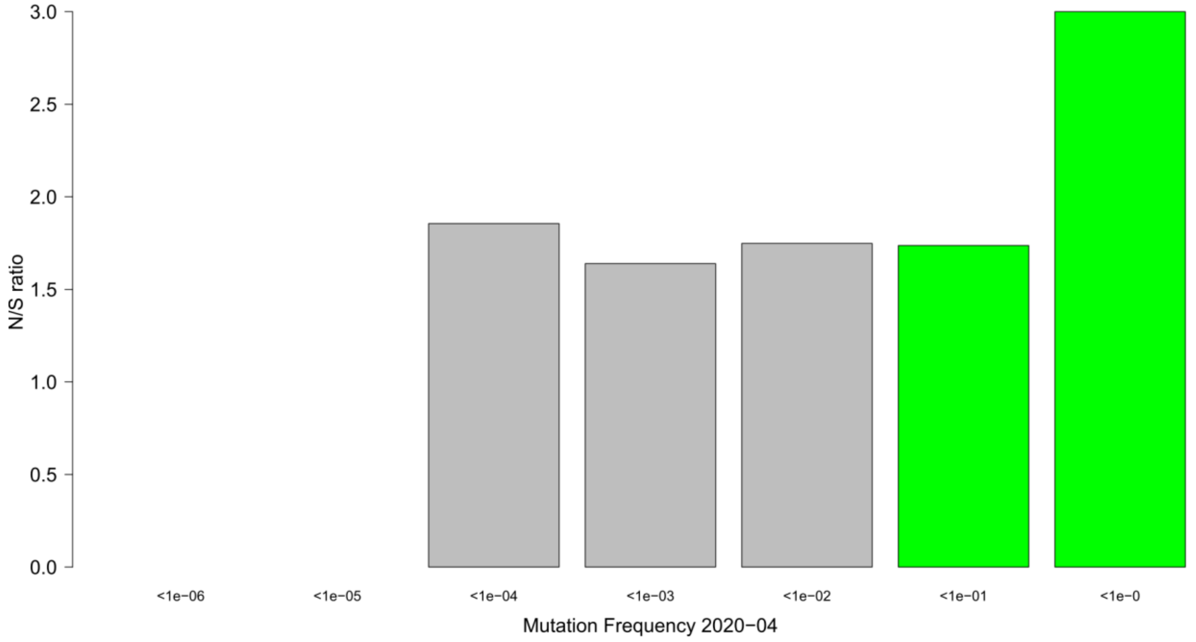

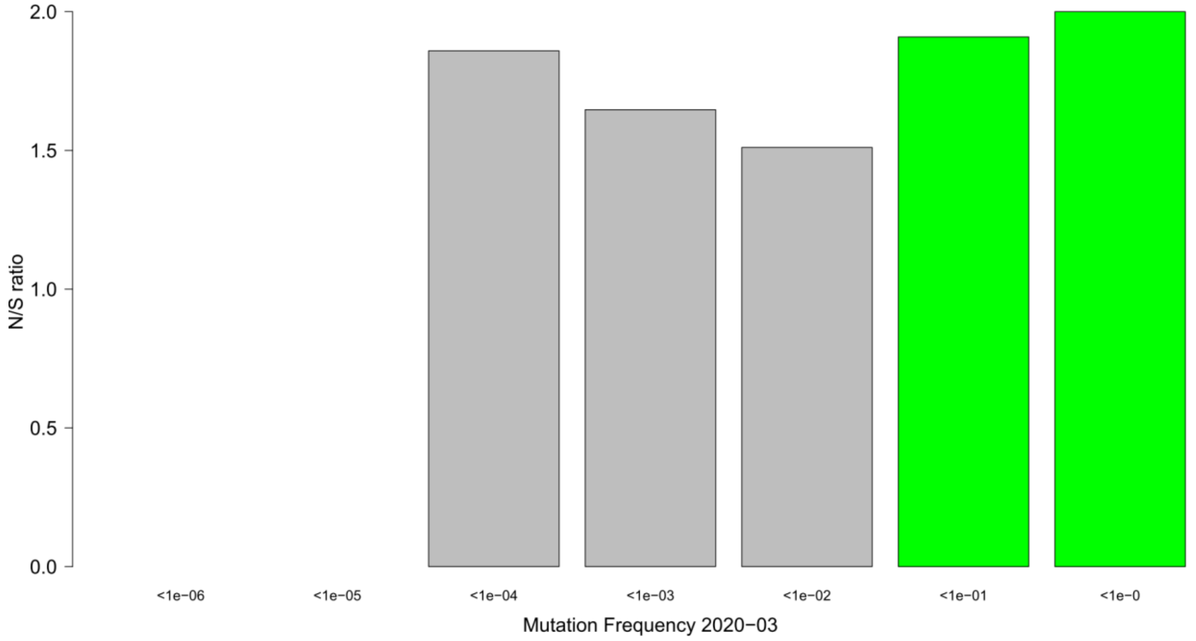

### Supplement Figure 4

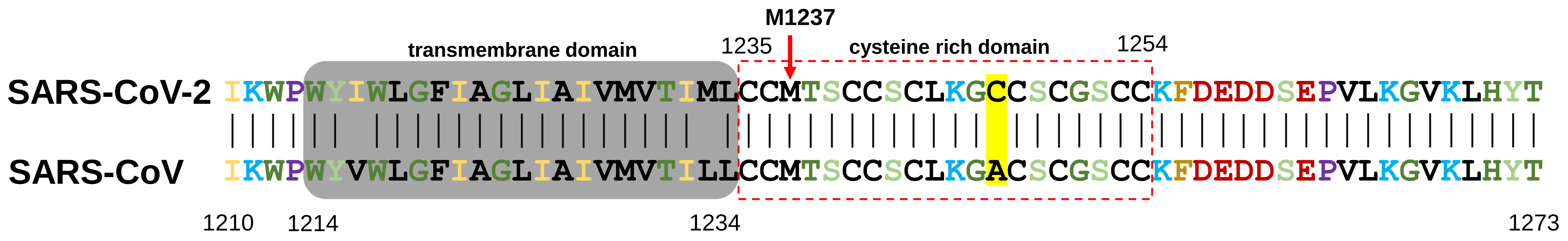


Fig. S4 The amino acid alignment of cytoplasmic tail of the SARS-CoV and SARS-CoV-2 Spike proteins.
